# Influence of *cis*-regulatory elements on expression divergence in human segmental duplications

**DOI:** 10.1101/2025.10.03.680410

**Authors:** Colin J. Shew, Gulhan Kaya, Sean P. McGinty, Megan Y. Dennis

## Abstract

Human-specific segmental duplications (HSDs) contain millions of base pairs of sequence unique to the human genome, including genes that shape neurodevelopment. Despite their young age (<6 million years), HSD genes exhibit widespread regulatory divergence, with paralog-specific expression patterns documented across a variety of tissues and cell types. Using long-read expression and epigenomic data, we show that human-specific paralogs tend to have lower activity than the shared, ancestral ones. To systematically characterize the *cis*-regulatory elements (CREs) within HSDs and understand patterns of regulatory change in recently evolved gene families, we conduct a massively parallel reporter assay of 7,760 human duplicated and chimpanzee orthologous sequences in lymphoblastoid (GM12878) and neuroblastoma (SH-SY5Y) cell lines. A large proportion (14–24%) of sequences exhibit differential activity relative to the chimpanzee ortholog (or between human paralogs), mostly with small fold-differences. Combining measured activity levels across all assayed sequences, predicted differences in *cis*-regulatory activity correlate with mRNA levels in SH-SY5Y. Differentially active CREs validated for *CHRFAM7A, HYDIN2*, and *SRGAP2C* may contribute to paralog-specific expression patterns and thereby to human-specific traits. While we identify some changes in CRE activity within duplicated regions, consideration of adjacent, unique sequences suggests a larger contribution from genome positional effects. In all, this work shows that functional divergence of duplicated CREs contributes moderately to regulatory divergence of HSD genes and uncovers enhancers that are candidate drivers of human-specific regulatory patterns.

## INTRODUCTION

Gene duplication is a major driver of evolutionary innovation, generating novel genetic material on which mutation and selection can act. Duplications are widespread, comprising a substantial proportion of genes across all domains of life, and may enable evolution of new traits by facilitating relaxed selection via genetic redundancy (Ohno 2013; Lynch and Conery 2000; Kondrashov et al. 2002). While a majority of paralogs are predicted to become pseudogenes and lost from the genome, the universal presence of gene duplications across species indicates that this process ultimately yields advantageous variation (Zhang 2003). Expression divergence is thought to be integral to gene retention post-duplication; the loss of *cis*-regulatory elements (CREs) can partition expression of daughter paralogs, eroding their redundancy and subjecting them to purifying selection (Force et al. 1999). Indeed, paralogs exhibiting expression divergence are predicted to persist, and may subsequently accrue additional regulatory or functional changes (Rodin and Riggs 2003). Further, gene regulation is highly plastic and itself a major driver of evolution. Alterations to spatiotemporal expression patterns are largely modular and preserve coding sequences, and so are less likely to be deleterious (Prud’homme et al. 2007).

In primates, segmental duplications (SDs) are large blocks of nearly identical sequence (>1 kilobase pair (kb), >90% similarity) that are especially enriched in African great apes (Bailey et al. 2001). They are typically interspersed hundreds of kilobases apart and co-occur with additional structural variation (Marques-Bonet et al. 2009; Lan and Pritchard 2016). Thus, SDs hold a strong potential to alter gene regulation by duplicating and translocating both genes and CREs. By contrast, tandem duplications are more common in other mammals but leave daughter genes in a similar regulatory environment (Marques-Bonet et al. 2009; Lan and Pritchard 2016). Human-specific segmental duplications (HSDs) are SDs unique to our species, which are consequently young (<6 million years) and highly similar (>99% nucleotide identity). In spite of this, genes within HSDs exhibit tissue-specific expression patterns across diverse primary tissues and cell lines, and a majority of HSD gene families display quantitative and spatial expression patterns specific to derived paralogs (Dennis et al. 2017; Florio et al. 2018). Some of these changes to gene regulation likely underlie human-specific traits, such as innovations in the development of the cerebral cortex. For example, human-specific *ARHGAP11B* drives basal neural progenitor proliferation and increased cortical neuron numbers in mammalian models (Florio et al. 2015; Kalebic et al. 2018; Heide et al. 2020), and is preferentially expressed in the germinal zone of the developing brain, while the ancestral *ARHGAP11A* is expressed at higher levels and more broadly throughout the neocortex (Florio et al. 2018). While *ARHGAP11B* has also attained novel biochemical functions (Namba et al. 2020), these differentiated expression patterns are also critical for human brain development.

The mechanisms underlying such cell-type specificity have not been thoroughly characterized. *Cis*-regulatory changes were broadly implicated in a study of HSD gene families that found reduced expression conservation in 5′-truncated genes, which had either lost their original or exapted novel promoters (Dougherty et al. 2018). Distal elements likely also contribute; we previously reported evidence for paralog-specific regulatory contributions from adjacent, non-duplicated genomic regions, as well as sequence-driven changes to the activity of duplicated enhancers (Shew et al. 2021). However, only a few gene families were functionally tested, and one of them (*ARHGAP11*) showed discordant mRNA levels and *cis*-regulatory activity between paralogs. This highlights the need to more comprehensively dissect the regulatory landscape of many HSD gene families, in order to gain mechanistic insight into how gene regulation diverges on short evolutionary timescales and might contribute to human-specific traits.

In this work, we assessed paralogous expression divergence in 605 human-duplicated gene families using a combination of short-read and long-read RNA sequencing in two cell lines: the lymphoblastoid cell line (LCL) GM12878 and neuroblastoma SH-SY5Y. We then used these cell lines to investigate *cis* regulation in a focal set of 26 HSD gene families by conducting a massively parallel reporter assay (MPRA) to directly compare human paralogous and chimpanzee orthologous CRE activity. We aimed to quantify functional divergence in these elements, link regulatory and expression differences between paralogs, and nominate CREs potentially contributing to human-specific expression patterns. We also used short- and long-read based histone modification information to consider the relative importance of duplicated CREs and adjacent, unique sequences outside of duplication breakpoints.

## RESULTS

### Human-duplicated genes are differentially expressed

Our previous work characterizing a subset of HSD genes identified divergent expression patterns over a short evolutionary time span (Shew et al. 2021). Leveraging recent work identifying additional duplicated gene families with at least one paralog unique to humans (Soto et al. 2025), we sought to expand our understanding of gene expression and regulation in HSDs. We also generated long-read RNA-seq data (PacBio Kinnex) in GM12878 and SH-SY5Y, two cell lines amenable to functional genomic assays. LCLs like GM12878 are accessible and directly comparable between primate species, and SH-SY5Y retains a neuronal progenitor-like state without *in vitro* reprogramming (Pezzini et al. 2017). Considering 605 duplicate gene families containing 1,394 genes, long- and short-read transcript abundance (Pezzini et al. 2017; ENCODE Project Consortium 2012; Luo et al. 2020; Hitz et al. 2023) (Supplemental Table S1) are strongly correlated in both cell types (*r*=0.85 for GM12878 and 0.88 for SH-SY5Y; Supplemental Fig. S1). While the long-read quantification is expected to be more accurate for distinguishing paralogs from each other, it is less sensitive overall, with a larger proportion of genes not detected in either cell type (17.5% with long reads versus 9.0% with short reads; Supplemental Table S2). Across duplicated gene families, we saw an enrichment of highest-expressed paralogs in regions with synteny to the chimpanzee ortholog (*p*=2.7 × 10^-4^ for GM12878 Kinnex; *p*=1.7 × 10^-3^ for GM12878 short-read; *p*=2.8 × 10^-6^ for SH-SY5Y Kinnex; *p*=2.6 × 10^-3^ for SH-SY5Y short-read) (Figure 1). This is in agreement with our previous work showing ancestral paralogs exhibit highest expression, as opposed to human-specific, derived genes (Shew et al. 2021). To the same end, we also found an enrichment of coding genes as opposed to pseudogenes among highest-expressed paralogs (*p*=2.7 × 10^-8^ for GM12878 Kinnex; *p*=1.5 × 10^-3^ for GM12878 short-read; *p*=2.0 × 10^-10^ for SH-SY5Y Kinnex; *p=*7.7 × 10^-3^ for SH-SY5Y short-read; Figure 1). This same enrichment was observed in RNA-seq data from primary adult liver, kidney, heart, and lung tissue, additional LCLs, and fetal neocortex (Florio et al. 2015; Fietz et al. 2012; Blake et al. 2020; Pickrell et al. 2010) (Supplemental Fig. S2). Taken together, these analyses confirm that human-specific genes exhibit divergent expression patterns, and the ancestral paralogs are most likely to retain a conserved function.

**Figure 1.**
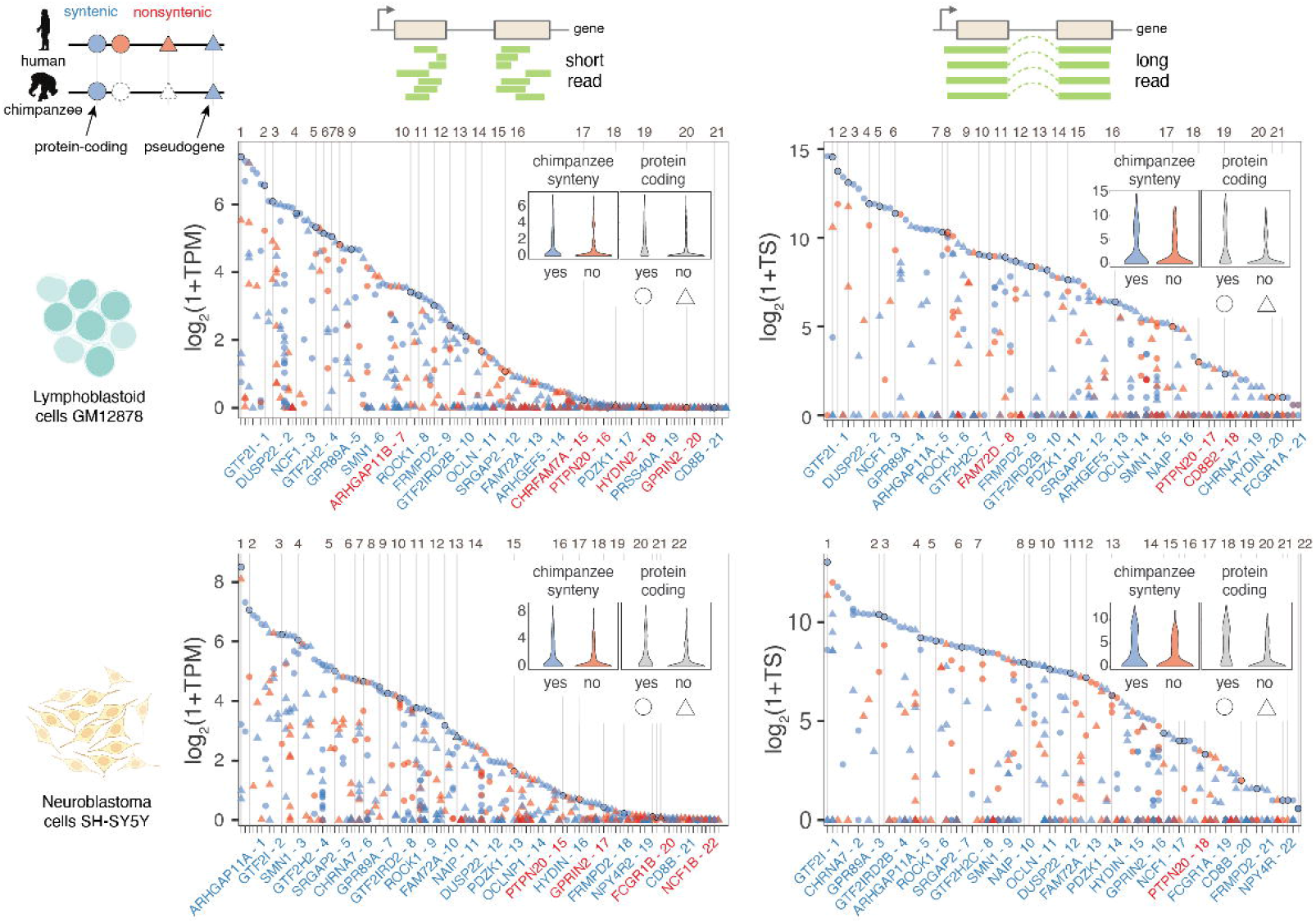
Human-duplicated genes are differentially expressed. The highest-expressed paralogs in each gene family (top point per column and inset violin plots) are more likely to be syntenic with chimpanzee (blue), and more likely to be protein-coding (circle). Comparisons were performed with transcriptomic data from GM12878 lymphoblastoid (top) and SH-SY5Y neuroblastoma cells (bottom) sequenced with short-read Illumina (left) and long-read PacBio Kinnex (right) reads. Gene families are represented on the x-axis, and transcript abundance on the y-axis (transcripts per million (TPM), TS (total transcript)). Gene family expression is ordered from highest (left) to lowest (right) with high-confidence human-specific duplications—which are the focus of the remainder of the study—numbered and demarcated with a vertical line. The corresponding number followed by the name of the highest-expressed paralog of these priority gene families is shown below the plot and colored to indicate syntenic status: likely ancestral (blue) or nonsyntenic, likely-derived (red).

### MPRA identifies active regulatory elements in human duplications

To systematically quantify *cis*-regulatory activity in HSDs, we designed an MPRA to directly compare paralogous human and orthologous chimpanzee sequences from 67 genes in 26 gene families (Figure 2A; Supplemental Tables S3, S4). These gene families were chosen based on their correct representation in the GRCh38 assembly (Nurk et al. 2022; Liao et al. 2023), known elevated copy numbers in all humans surveyed compared with nonhuman primates (Soto et al. 2025), and previously established ancestral and derived identities (Dennis et al. 2017). Additionally, our expression analysis shows 72% (18/25) of these gene families detected with short reads (>1 TPM) in GM12878 or SH-SY5Y, and 68% (17/25) with long reads (>1 CPM) (Supplemental Table S5). We used chromatin accessibility and histone H3, lysine 27 acetylation (H3K27ac) chromatin immunoprecipitation sequencing (ChIP-seq) data to identify candidate CREs from fetal and adult prefrontal cortex (Vermunt et al. 2016; Reilly et al. 2015; de la Torre-Ubieta et al. 2018)—due to the known role of HSD genes in brain development—and LCLs (McVicker et al. 2013; Degner et al. 2012; Bryois et al. 2018)—whose accessibility has made them foundational for cross-species comparisons in primates (Supplemental Table S1). HSDs are typically excluded by standard bioinformatic approaches, as reads aligning to these regions have poor mapping quality. To circumvent this, we permissively identified candidate CREs by re-mapping epigenomic data to the human reference (GRCh38) with multiple alignments. A total of 1,163 CREs in ancestral loci were divided into 200mer “tiles” with 100-bp overlap (N=2,675), and homologous tiles in the human and chimpanzee (panTro6) genomes were identified by sequence alignment. In total, we synthesized 8,145 test sequences and 500 scrambled negative controls (Supplemental Fig. S3, Supplemental Table S6), which were packaged into lentiviral vectors and assayed in GM12878 and SH-SY5Y cells according to the lentiMPRA protocol (Gordon et al. 2020). While these are cancer-derived cell lines with abnormal karyotypes, they provide a relevant *trans*-regulatory environment, and CRE activity is measured at many genomic positions. Transduction efficiencies were comparable to previous lentiMPRA studies, with multiplicity of infection (MOI) of ∼40 for SH-SY5Y and ∼13 for GM12878. Technical replicates (N=3 per cell line) exhibited high reproducibility (mean pairwise r=0.93 and 0.93 for within-cell type comparisons of DNA and RNA libraries, respectively; Supplemental Fig. S4) (Liao et al. 2023; Nurk et al. 2022).

**Figure 2.**
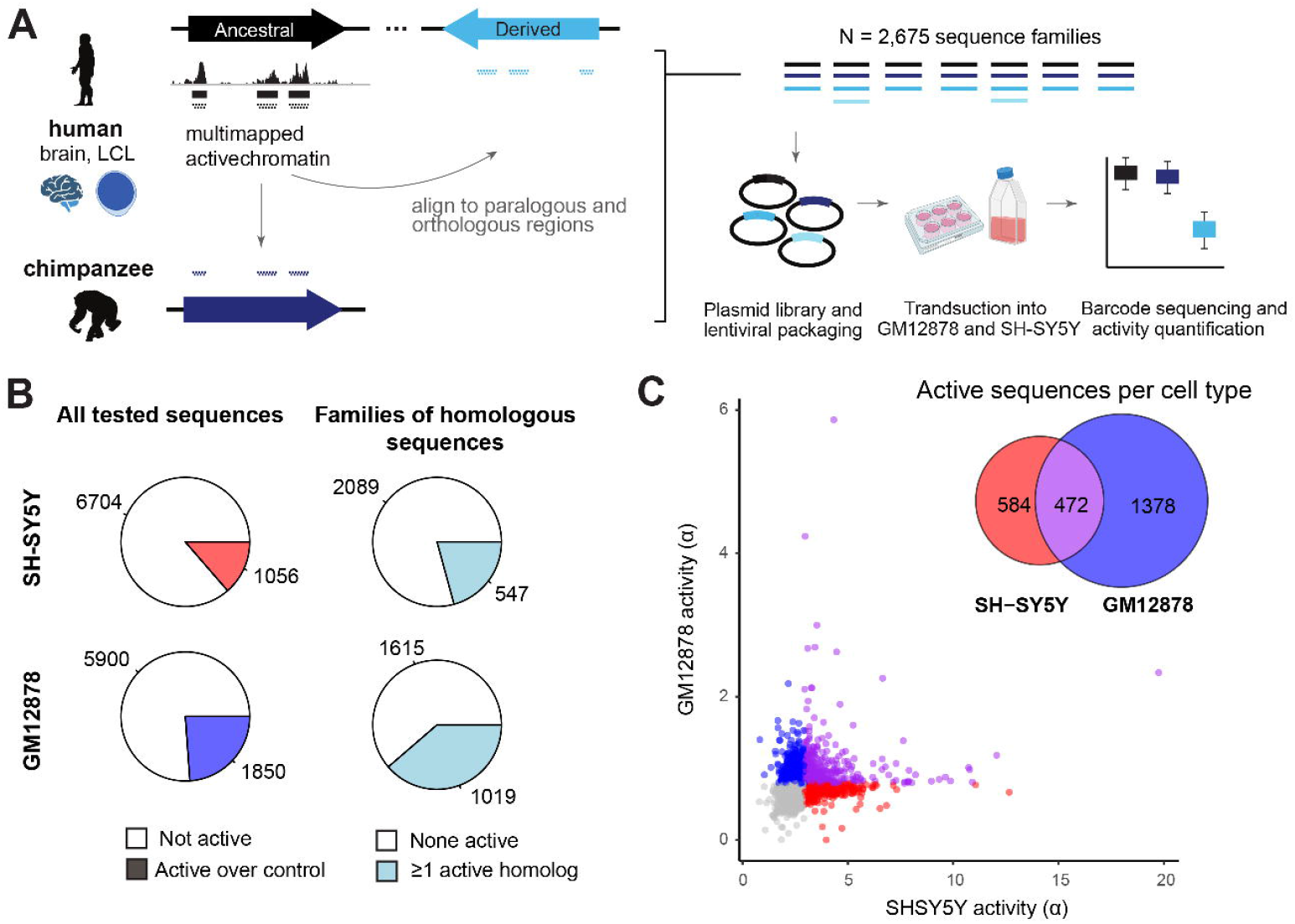
MPRA quantifies paralog-specific regulatory activity of duplicated CREs. **(A)** MPRA tiles were designed to multi-mapped chromatin accessibility and histone H3K27ac ChIP-seq data previously published from human primary brain (fetal and adult) and lymphoblastoid cell lines. Peak regions were tiled with 200mers at 2x density in ancestral loci, and homologous positions were identified by alignment to the human and chimpanzee genomes. All unique sequences were synthesized in an oligonucleotide pool and cloned into a lentiviral vector. MPRA was performed in the neuroblastoma cell line SH-SY5Y and lymphoblastoid cell line GM12878. **(B)** Pie charts depict the number and proportion of active sequences (left) and sequence families containing at least one active sequence (right), for each cell type. **(C)** Sequence activity (estimated transcription rate, □) is plotted type for 7,750 tiles measured in both assays. Sequences active over negative controls are colored red (SH-SY5Y), blue (GM12878), and purple (both). The Venn diagram indicates the number of sequences in each category.

We first quantified enhancer activity for all tested CREs in both cell lines using the estimated transcription rate (□), modeled from the integrated (DNA) and expressed (RNA) barcode counts (MPRAnalyze (Ashuach et al. 2019)). In SH-SY5Y, 1,056/7,760 (14%) tested sequences were active relative to the negative controls (median absolute deviation *p*<0.05), while in GM12878, 1,850/7,750 (24%) sequences were active (Figure 2C; Supplemental Tables S7–S8). Of these, 472 (19%) of 2,434 active sequences were active in both cell types (Figure 2B; Supplemental Note S1). Notably, higher activity was measured overall in SH-SY5Y (median □=2.50, versus 0.73 in GM12878; Figure 2C), likely due to the three-fold greater infection rate achieved in this cell line. Activity measurements were correlated for overlapping tiles (*r*=0.21 for GM12878 and 0.43 for SH-SY5Y; Supplemental Fig. S5C), but we opted against combining measurements across tiles because any target nucleotide was covered by at most two constructs, and the 100-bp offset is much larger than transcription factor binding sites.

Sequences within 1 kb of transcription start sites (TSSs) were enriched for activity in both cell types (1.35-fold higher proportion of active sequences for promoters in GM12878; 3.46-fold in SH-SY5Y; Fisher’s exact test *p*<0.02). While we observed a correlation between promoter activity (mean activity score for all active tiles) and mRNA expression driven by the contrast between active and inactive promoters (Supplemental Fig. S7A), MPRA promoter measurements did not predict differences in ancestral/derived gene pairs (Supplemental Fig. S7B). This suggests a role for distal elements in shaping expression patterns.

### Species- and paralog-specific CRE activity

We first examined sequence activity across species. Considering all human-chimpanzee sequence pairs with at least one active element, 35% (357/1,017) were differentially active in SH-SY5Y, and 11% (220/1,935) were differentially active in GM12878 (Supplemental Fig. S8; Supplemental Tables S9 and S10). We found an expected relationship between sequence divergence and the magnitude of differential activity between species in both cell types (*r*=-0.07, *p*<1×10^-3^; Supplemental Fig. S9). However, the strongest differentially active sequences tended to have the highest similarity with chimpanzee, demonstrating that a small number of substitutions can alter sequence activity (Supplemental Fig. S9). Based on our findings that ancestral genes generally exhibit higher expression and increased concordance with chimpanzee orthologs (Shew et al. 2021) (Figure 1), we next hypothesized that activity losses preferentially occur in derived CREs. We partitioned human sequences by ancestral or derived status, but found no enrichment of activity gains or losses (considering chimpanzee a proxy for ancestral activity) in either group (hypergeometric test). However, the measured activities of ancestral sequences correlated significantly better than derived sequences with their chimpanzee orthologs (Fisher’s z-test, *p*<0.01 in both cell types; Figure 3A). Thus, while *cis*-regulatory activity in HSDs is largely conserved between humans and chimpanzees, human-specific derived regions are more diverged, potentially experiencing relaxed evolutionary constraint on CRE activity.

**Figure 3.**
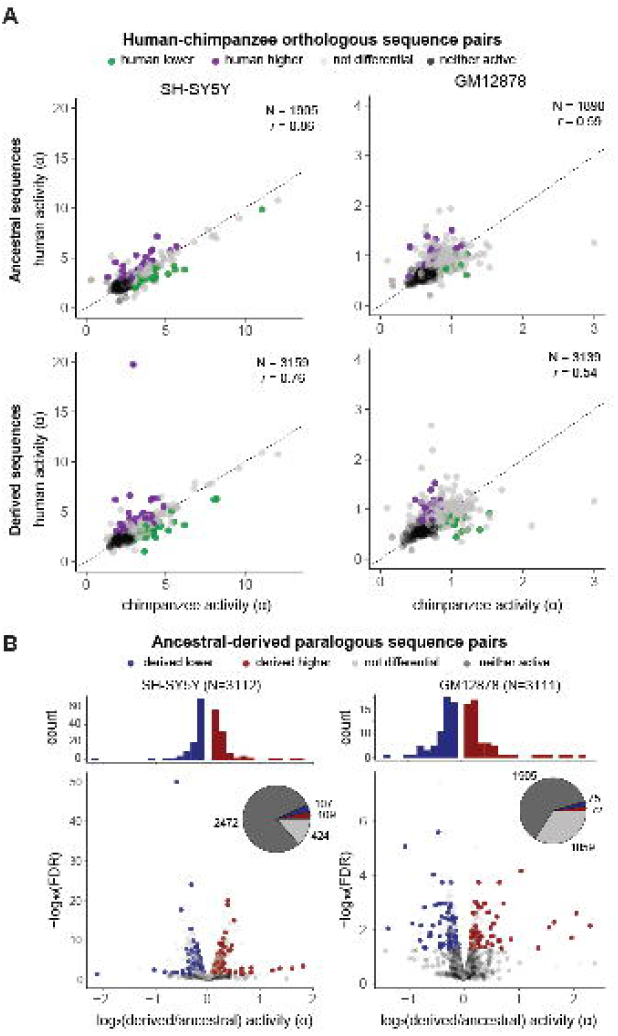
Differential activity between homologous sequences. **(A)** Scatterplots depict the activity of human and orthologous chimpanzee sequences, partitioned by human evolutionary status (ancestral above, derived below). Each sequence pair is colored by differential activity relative to chimpanzee: green (human lower), purple (human higher), gray (not differential), black (neither sequence active). The dotted line marks the identity y=x, and Pearson’s correlation coefficient is printed on each plot. **(B)** Volcano plots depict differential activity between paralogous human sequences, with each unique derived-ancestral pair plotted separately. Points are colored by differential activity relative to the ancestral human paralog: blue (derived lower), red (derived higher), gray (not differential), black (neither sequence active). The marginal histograms depict the distribution of log-fold differences for differentially active sequences.

Comparing activities between human duplications, we found a similar proportion of derived-ancestral pairs were differentially active by the same definition: 216/640 (34%) of pairs in SH-SY5Y, with a median 1.13-fold difference; in GM12878, 147/1,206 (12%) were differentially active, with a median 1.20-fold difference (Figure 3B–C). While most paralogs exhibited modest divergence in activity (1.13-fold and 1.20-fold differences in SH-SY5Y and GM12878, respectively), *SRGAP2* and *FRMPD2* contained strong differential human-specific sequences (>2-fold; Supplemental Tables S11 and S12) that were also concordant in both cell types, suggesting possible single CRE drivers of paralog differences. Targeting the putative CRE from *SRGAP2* (Chr 1:206333093-206333293; GRCh38), we used a luciferase reporter assay to validate that a larger homologous region containing the human-specific *SRGAP2B* (0.43-fold) and *SRGAPC* (0.36-fold) tile sequences were significantly less active than the ancestral *SRGAP2* in SH-SY5Y, with no differential activity observed in GM12878, matching MPRA results (Supplemental Fig. S10; Supplemental Table S13).

Taking this same approach to test larger regions with luciferase reporters, we used 1-kb windows in HSDs to score total measured MPRA activity, secondarily ranking by the maximum fold difference of any contained tile, and manually curating select differential elements in both cell types (Supplemental Fig. S11). In the *CHRNA7* locus, we tested two candidate regions. The first (Chr 15:32183316-32184316; GRCh38) was predicted to have higher activity in the derived *CHRFAM7A*, which was true for GM12878 (>1.3-fold difference; Figure 4 and Supplemental Figure S12; Supplemental Table S13). The second was expected to have lower activity in SH-SY5Y, which was also verified by luciferase (<0.7-fold difference). In addition, an intronic region within ancestral *HYDIN* was predicted to have higher activity in the derived *HYDIN2* in both cell types. Reporter activity was significantly higher for both homologs in GM12878, and for *HYDIN2* in SH-SY5Y (>3.8-fold difference; Supplemental Fig. S12 and S13; Supplemental Table S13). Altogether, these experiments demonstrate paralog-specific activity for enhancers in a more biologically-relevant-sequence context. The reporter activities from these constructs comprising expanded putative CREs (1 kb) were generally in agreement with the MPRA, which measured activity of 200mer tiles in isolation.

**Figure 4.**
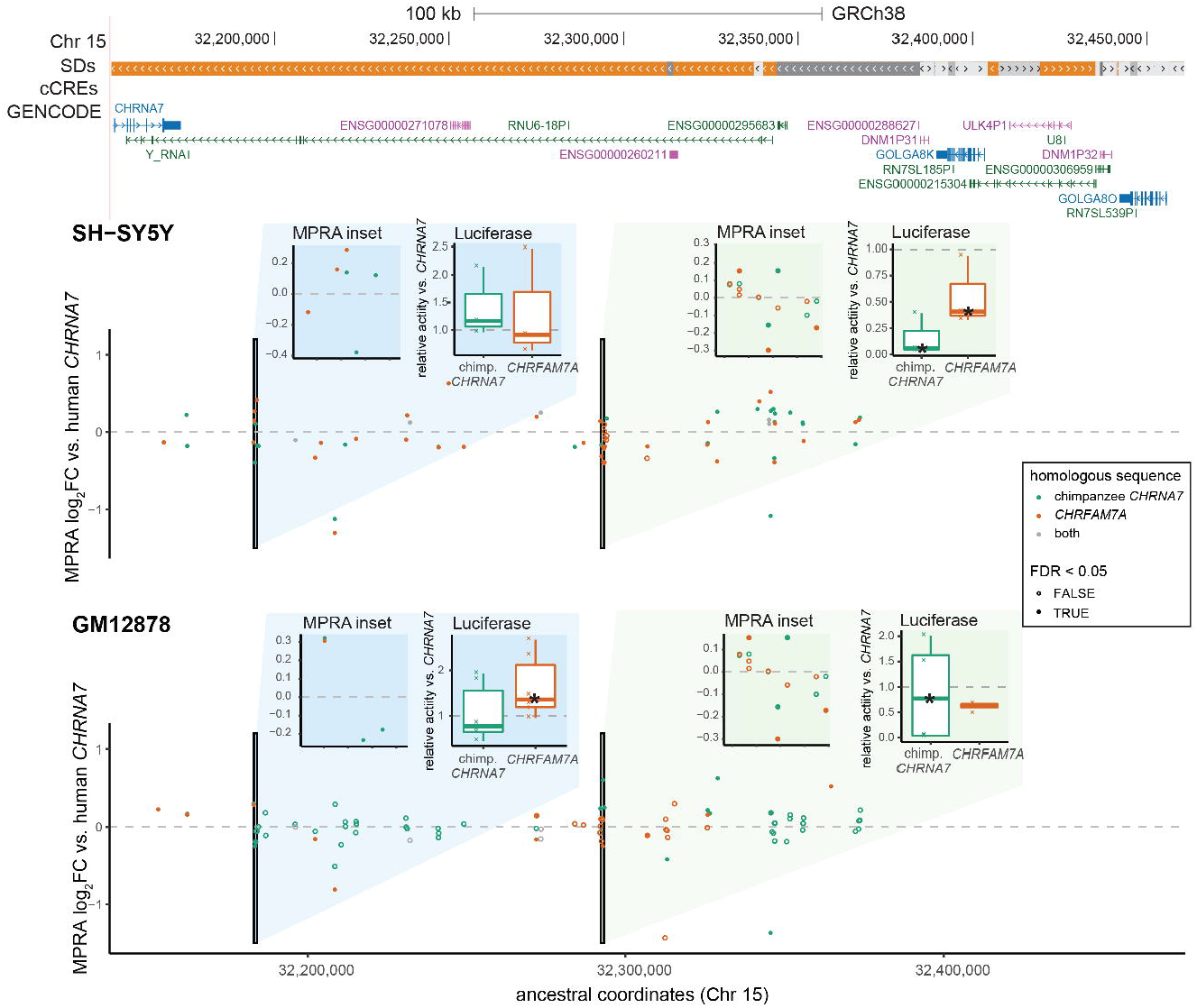
Paralog-specific regulation in the *CHRNA7* locus (Chr 15q13.3). A 143-kb region (Chr 15:32,153,206-32,460,660; GRCh38) is shown. Segmental duplications (SDs) are colored orange for >99% sequence identity, and gray for <98% identity. No ENCODE candidate *cis*-regulatory elements (cCREs) are annotated in this locus. The MPRA data are visualized as activity relative to the human ancestral locus, and plotted on these coordinates. Each point represents a 200mer tile, and tiles are color by which homolog they correspond to (teal: chimpanzee *CHRNA7*; orange: human-specific *CHRFAM7A*; gray: identical in both) and filled in if they scored as differentially active. Two 1-kb regions are highlighted in the insets; the left panel shows a zoomed in plot of the MPRA measurements per tile, and the right shows a relative luciferase activity of the entire sequence, as assayed in the corresponding cell type. Asterisks on the box plots mark significantly different activity with respect to the human ancestral sequence. See also Supplemental Table S13.

### Collective analysis of CREs on paralog expression

Due to modest effect sizes of individually tested CREs in the MPRA, we next considered whether their collective activity might explain paralog-expression divergence. Building on the activity-by-contact (ABC) score (Fulco et al. 2019), we developed a similar metric using activity levels directly measured from the MPRA (Methods; Supplemental Fig. S14A). For expressed derived-ancestral gene pairs (either TPM>1), expression divergence in the short-read data was significantly correlated with the summed ABC score ratio in SH-SY5Y (p<0.05, r=0.70; Supplemental Fig. S14B). This comparison was robust to various cutoffs for considering CREs active or differential and showed the best correlation for more stringent criteria. No relationship was observed using the long-read expression data, however, perhaps reflecting reduced sensitivity of Kinnex to detect transcripts (Supplemental Fig. S14C; Supplemental Note S2). The SH-SY5Y MPRA ABC score ratio also correlated with paralog expression divergence quantified in certain primary brain cell types (i.e., the cortical plate, cortical neurons without basal contact, and the outer subventricular zone; Supplemental Fig. S16), reflecting functional relevance beyond the cell lines assayed.

We searched for trends in CRE activity that could explain this, namely in strength, number, or location. In both cell types, neither measured CRE activity nor the number of CREs differed between ancestral and derived regions, but active CREs were slightly closer proximally to ancestral TSSs than derived ones (*p*=0.03, Wilcoxon rank-sum test; Supplemental Fig. S17). This suggests a trend toward loss of active elements closer to derived paralogs. Because promoter strength alone did not predict expression differences (Supplemental Fig. S7), considering the full complement of CREs may better explain differential mRNA levels across HSD genes.

Indeed, the fraction of contribution of proximal CREs (within 1 kb) as determined by the ABC metric was generally concordant across paralogs (Supplemental Fig. S18A). There were a few exceptions, namely *ARHGEF34P* and *ARHGAP11B* in SH-SY5Y and *PTPN20CP* and *HYDIN2* in GM12878 (Supplemental Fig. S18B–C). Across all elements, the median fraction of contribution of the top CRE to each HSD gene was only 5% (Supplemental Fig. S18D). Together, these results indicate that changes to promoter activity are not the primary driver of HSD expression divergence in most gene families, and that many distal elements may influence this process.

For comparison, we also calculated the predicted difference using the ABC score as published, with chromatin accessibility and H3K27ac ChIP-seq. We aligned datasets from GM12878 (Zhang et al. 2020) and SH-SY5Y (Gartlgruber et al. 2021; Zimmerman et al. 2018) to the human reference (GRCh38), allocating multiple alignments probabilistically (Zhang et al. 2020; Chung et al. 2014), enabling the calculation of the ABC difference metric across HSDs and adjacent, unique regions (Methods). For SH-SY5Y, we found a correlation of the ABC score ratio with short-read expression divergence (*p*=0.05, *r*=0.42), but this did not hold when subsetting to only the duplicated or unique space (Figure 5A–C). The long-read expression data for both cell types also trended in the expected direction for all and non-duplicated peaks. We also scored the exact regions tested by our MPRA, in place of peaks derived from the epigenomic data, and found a weaker correlation than the MPRA analysis on its own (*p*=0.06, *r*=0.39) (Figure 5D). Thus, the measured regulatory activity from the MPRA appears more informative than proxies based on chromatin state. The whole-genome chromatin activity data, however, suggest that more distal, non-duplicated CREs have a greater impact on expression divergence.

**Figure 5.**
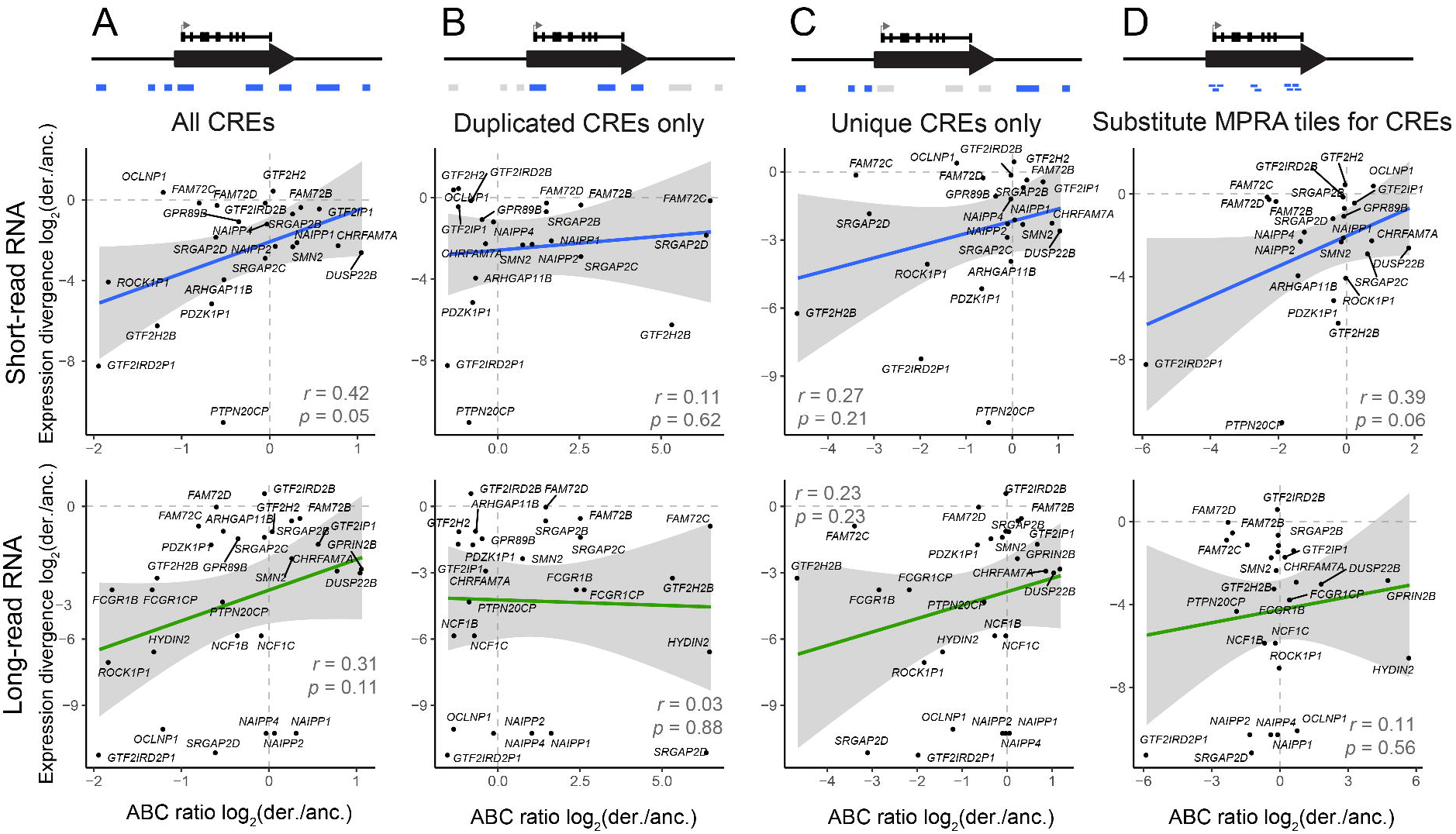
Comparison of SH-SY5Y expression divergence to predictions from epigenomic data. For each derived-ancestral (der./anc.) gene pair, expression divergence was correlated to the ratio of summed ABC scores calculated from multi-mapped ATAC-seq and H3K27ac ChIP-seq. The comparison was performed on the following sets of regions: **(A)** all CREs within 5 Mb, as defined by our ChIP-seq analysis; **(B)** CREs within HSDs; **(C)** CREs outside of HSDs; or **(D)** the exact positions tested by MPRA. Pearson’s correlation and significance of the linear relationship is printed on each plot, and the 95% confidence interval for the regression is shaded in gray.

Finally, recent advances in long-read epigenomic assays have better enabled differentiation of regulatory information between human duplications (Gershman et al. 2022; Altemose et al. 2022). Using published directed methylation with long-read sequencing (DiMeLo-seq) datasets from GM12878 (Maslan et al. 2024), we identified expected relationships between DNA methylation (5mC), histone modifications (H3K4me3, H3K27ac) and HSD gene expression, including derived-ancestral differences (Supplemental Fig. S20A). These data corroborated our findings using short-read ChIP, identifying a similar correlation between regulatory activity (ABC score) and expression divergence that was primarily driven by non-duplicated elements (Supplemental Note S3). Moving forward, we expect long-read epigenomic datasets will be critical for disentangling the mechanisms underlying paralogous expression divergence in other loci and cell types.

## DISCUSSION

In this study, we present the first high-throughput quantification of regulatory activity in duplicated regions. Broadly, HSD sequences exhibit few strong differences in CRE activity. This is unsurprising given the small number of substitutions between human paralogous and chimpanzee orthologous sequences, and comparable to the fraction of differentially active sequences seen in MPRAs of polymorphic and species-specific variants (Uebbing et al. 2021; Tewhey et al. 2016). However, this stands in contrast to examples of multi-fold differences of HSD promoter and enhancer activity demonstrated previously (Shew et al. 2021). One explanation may be the size of assayed sequences; MPRA libraries constructed from synthetic fragments are currently limited to ∼200 base pair insert sizes, while previously tested CREs in HSDs assayed fragments >1000 base pairs in size. Indeed, one of the regions we tested (*HYDIN*) in a luciferase assay displayed an almost four-fold difference in activity, while individual tiles measured below two-fold difference by MPRA (Supplemental Fig. S13). It is therefore likely that the synergistic effects of nearby enhancers are not completely captured in the tiled oligonucleotide design.

Despite these limitations, MPRA is well suited for directly comparing homologous sequences. Assessing CRE activity between humans and chimpanzees, we found that derived sequences were more weakly correlated than those from ancestral paralogs. This is in line with the predicted relaxation of selection on derived genes, though gains and losses of activity were seen in similar proportion. In order to account for both the strength and distance of duplicated elements across human paralogs, we used MRPA measurements to develop a metric based on the ABC score. This integrated score predicted expression differences, albeit only using the short-read RNA quantification. Meanwhile, consideration of a larger complement of CREs using chromatin-based proxies for activity suggests adjacent, non-paralogous regions are collectively more impactful. Overall, these results paint a picture in which sequence divergence of human-duplicated regulatory elements influences but does not primarily drive changes to expression of paralogous genes.

We also highlighted individual differentially active elements that may contribute to paralog-specific gene regulation. In the *CHRNA7* locus, we tested two regions containing multiple differentially active MPRA tiles: the first was significantly more active in *CHRFAM7A*, and the second was less active in SH-SY5Y, tracking with RNA expression and CRE activity in the MPRA. *CHRNA7* codes for the cholinergic receptor nicotinic alpha 7 subunit and resides in a genomically unstable region in Chromosome 15q11-q13 where copy-number variants are associated with neurodevelopmental features, including intellectual disability, autism, and epilepsy (Watson et al. 2014). Human-specific *CHRFAM7A*, meanwhile, is a partial duplication and fusion gene linked to schizophrenia and shown to interact with *CHRNA7* as a dominant negative (Gillentine et al. 2017). Accordingly, expression modulation is likely critical to proper dosage. If these CREs indeed regulate *CHRNA7*, the second may play a role in reducing *CHRFAM7A* expression in the brain, as indicated by its lower activity in SH-SY5Y. Understanding the cell type-specific activity of these and other CREs will be critical to unraveling their function.

This study focused on comparisons of CRE activities, leaving the contributions from changes in chromatin conformation at HSD and orthologous chimpanzee loci an open question. Identification of promoter-enhancer loops would enable more confident assignment of CREs to target genes, identification of paralog-specific chromatin contacts across duplication breakpoints, and refinement of ABCD regulatory predictions by using true interaction frequency in place of a genomic distance proxy. While published methods for allocation of multimapping Hi-C reads exist, they still exhibit reduced performance in highly identical HSD regions (Zheng et al. 2019). Long-read based chromatin conformation capture methods such as Pore-C (Deshpande et al. 2022) and CiFi (McGinty et al. 2025), with larger fragment sizes, will be necessary to confidently identify loops involving HSD regions. Finally, machine learning methods are rapidly improving prediction of epigenetic properties from DNA sequence alone.

Chromosome conformation, accessibility, histone methylation, and RNA expression can all be compared *in silico*, though accurate variant effect prediction is currently an area of active development, and comparisons between nearly identical HSD paralogs would also require validation. The proliferation of telomere-to-telomere genomes also raises the possibility of expanding analyses to additional recent gene duplications now resolved in these assemblies (Nurk et al. 2022; Vollger et al. 2022), as well as differentiating allelic and paralogous single-nucleotide variants within HSDs and orthologs to expand on our study of a single haplotype per species (Supplemental Note S4).

In all, this work measured the regulatory activity of thousands of candidate regulatory sequences distinguishing recent human duplications. We found that a large proportion of them are active, and while few individual 200mers exhibit strong differential activity, some contribute in combination to regulatory patterns of HSD genes such as *CHRFAM7A, HYDIN2*, and *SRGAP2C*, perhaps driving unique features of our species. Integration of additional information, such as new long-read epigenomic datasets, chromatin conformation, and single nucleotide and structural polymorphism will be necessary to gain a more complete picture of gene regulation within HSDs.

## MATERIALS AND METHODS

### RNA library preparation

Total RNA was extracted from GM12878 and SH-SY5Y cell lines using the Qiagen RNeasy Plus Kit following the manufacturer’s protocol. RNA integrity was confirmed using the Agilent 2100 Bioanalyzer, ensuring RIN values ≥7.0. Full-length RNA libraries were prepared using the PacBio Kinnex Full-Length RNA Kit (103-238-700), including cDNA synthesis, Kinnex PCR amplification, and library cleanup. Libraries were quantified with the Qubit DNA HS Assay Kit and size-checked on the Agilent Femto Pulse system, which showed average insert sizes of ∼15 kb for both GM12878 and SH-SY5Y, before sequencing on the PacBio Revio platform.

### RNA-seq analysis

RNA-seq data were obtained for GM12878 (ENCODE ENCSR000AEC, ENCSR000AEE, and ENCSR000CVT) and SH-SY5Y (Pezzini et al. 2017). Transcripts were quantified with Salmon v1.9.0 (Patro et al. 2017) with the flags “--validateMappings --gcBias”, the telomere-to-telomere CHM13 v2.0 CAT/Liftoff transcriptome, and the CHM13 v2.0 assembly as decoy sequence. All identical transcripts were removed from the transcriptome prior to index construction. Transcripts per million (TPM) values were summed to the gene level using tximport (Soneson et al. 2015). For relative expression analyses, derived-ancestral gene pairs were considered if at least one paralog was expressed >1 TPM. Expression divergence was calculated as the log_2_ ratio of derived/ancestral expression, with a pseudocount one order of magnitude smaller than the smallest nonzero value added to each. For direct comparison of count-normalized datasets, raw reads were downsampled with seqtk v 1.4.

### Long-read PacBio Kinnex analysis

Initial processing of PacBio Kinnex data was done with the Iso-Seq workflow (https://isoseq.how/). HiFi reads were first segmented using PacBio concatenated read splitter Skera at adapter positions to generate segmented reads. Lima is then used to identify and remove barcoded Iso-Seq primers. The output of lima are full length (FL) reads. These FL reads are then processed with Iso-Seq refine to identify and trim poly(A) tails, as well as identify and remove concatemers. The output of Iso-Seq refinement is full-length, non-concatemer (FLNC) reads. These FLNC reads are then quantified using Oarfish (Zare Jousheghani et al. 2025) using the same identical transcript-filtered CHM13 v2.0 CAT/Liftoff transcriptome described above. The estimated transcript counts are then summed to get gene-level counts, which were converted to counts per million (CPM) for expression analyses. No length normalization was performed because each transcript detected is expected to yield one FLNC read.

### MPRA oligo design

Candidate CREs were identified from H3K27ac ChIP-seq, ATAC-seq, or DNase-seq data in LCLs, fetal cortex, and adult prefrontal cortex with data from the following publications: (McVicker et al. 2013; Degner et al. 2012; Bryois et al. 2018; Vermunt et al. 2016; Reilly et al. 2015; de la Torre-Ubieta et al. 2018). In the case of Bryois et al., only control samples were used. Reads were trimmed with Trimmomatic (Bolger et al. 2014) aligned to GRCh38 allowing for multiple mapping with Bowtie v1.1.2 (Langmead et al. 2015) with the flags “-a -v2 -m 99”. Peaks were called using MACS2 v2.1.2 (Liu 2014) with a 5% FDR and the following additional settings for chromatin accessibility data: “--nomodel --shift - 100 --extsize 200 --broad”. Peaks were called on all samples in each study and combined by reporting genomic regions with nonzero coverage in a minimum number of samples using BEDTools genomecov (Quinlan and Hall 2010): 3/10 LCL H3K27ac, 52/204 LCL DNase, 2/3 fetal cortex H3K27ac, 3/3 fetal ATAC-seq (cortical plate or germinal zone), 1/1 adult prefrontal cortex H3K27ac (peaks called jointly from samples HS1 and HS2), and 102/137 adult prefrontal cortex ATAC-seq. Cutoffs were chosen based on manual inspection of reproducibility and to equalize sequences represented from LCLs and brain.

To design the test oligos, all peaks within 400 bp were merged, the maximum distance at which peaks could become adjacent after coordinate expansion. The resulting intervals were padded to the nearest 100 bp, plus an additional 100 bp on either side to allow full 2x coverage of the initial target regions using 200mer tiles. Tiles were generated with BEDTools make windows. To match orientation of promoters, tiles were assigned to the strand of the nearest annotated feature (GENCODE v32), or randomly selected if multiple features were nearest. This resulted in random strand assignments to 140/661 promoter and 501/7,099 distal tiles. Tiles were intersected with ancestral HSD regions and aligned with BLAT (Kent 2002) to a custom reference consisting of GRCh38 with non-HSD sequence masked, in addition to the missing portions of the DUSP22B and GPRIN2B contigs (ancestral loci and contigs from (Dennis et al. 2017)). Alignments were then filtered in a two-step process to keep only paralogous positions found in human-specific segmental duplications: (1) keep if there were ≤4 hits >190 bp with >95% identity; (2) else keep if there were ≤4 hits >195 bp with >98% identity. This logic maximized target regions by allowing greater divergence for individual tiles on the first pass but increasing stringency if the number of hits was still greater than the expected number of paralogs. To focus on paralogous differences, sequences for which all alignments were identical were discarded. To identify chimpanzee orthologs, human tiles were mapped to the panTro6 assembly using liftOver (UCSC Genome Browser utilities) and filtered for being with 5% (10 bp) of the original 200 bp size (93% success rate). All tiles with this limit were trimmed or expanded to 200 bp; 91% of alignments were within 1 bp difference. FASTA sequences were extracted in a strand-aware manner from GRCh38 or panTro6 and deduplicated. Finally, a universal priming sequence (5’-AGGACCGGATCAACT-[200mer]-CATTGCGTGAACCGA-3’) to the pLS-SceI vector (Addgene 137725) was added to all 200mers. Sequences were filtered to remove AgeI and SceI restriction sites, as well as those made singletons after filtering, leaving 8,145 test oligos. In addition, 500 ancestral tiles were sampled and scrambled to generate negative controls. All 230mer sequences were synthesized by Agilent Technologies.

### MPRA library preparation and sequencing

MPRA was carried out according to the lentiMPRA protocol as described (Gordon et al. 2020) with 3 technical replicates. We generated the lentivirus library by transfecting HEK293T cells with the plasmid library. To titrate the lentivirus, GM12878 and SH-SY5Y were plated into a 24-well plate and infected with varying volumes of the virus in each well. In these experiments, transduction efficiency was ∼10% for GM12878 and ∼30–40% for SH-SY5Y cells, as determined by GFP-positive cell counts on the Countess II FL. Following a 3-day incubation, genomic DNA was extracted from each well. Then, the MOI was calculated for each viral concentration via qPCR using plasmid backbone and genomic DNA control primers, as described in the protocol. From these titration results, the final infection conditions were set as follows: For SH-SY5Y experiments, 2.5 million cells were infected at a multiplicity of infection (MOI) of 40 in DMEM/F12 Glutamax medium (Thermo Fisher Scientific 10565018) with 10% FBS (Thermo Fisher Scientific, 10-438-026) and 2.5 µg/mL protamine sulfate (Spectrum chemical TCI-P0675-1G). For GM12878, 55 million cells were transduced at an MOI of 13 in RPMI (Fisher Scientific 11875135) with 10% FBS and 8 µg/mL polybrene (Millipore Sigma MILL-TR-1003-G). Cells were incubated at 37°C in 5% CO_2_ and nucleic acid extractions were conducted 48 hours after infection. Barcode association was performed on a PE150 NextSeq mid-output run. Barcode counting from DNA and RNA libraries was performed on three PE15 NextSeq high-output runs.

### MPRA data analysis

Barcode-insert association was performed with MPRAflow (Gordon et al. 2020) using “--mapq 1” and all other parameters set to default. We manually confirmed that promiscuous barcodes had a majority assignment to one insert and were not mixed between paralogs. Barcodes were counted with the count utility and passed to MPRAnalyze (Ashuach et al. 2019). Counts were depth-normalized with the total sum, and active sequences were defined per cell type against the negative controls with the following model: “dnaDesign = ∼ barcode + replicate, rnaDesign = ∼ 1”. To identify differential activity of homologous sequences, counts matrices were reformatted to combine all homologs in the same row for model fitting, to use homolog identity used as a covariate. Models were fit in “scale” mode with the following design: “rnaDesign = ∼homolog, reducedDesign = ∼1”. The alpha value used as a measure of sequence activity refers to the estimated transcription rate from the model. Differentially active homologs were defined in two ways: (1) against the chimpanzee ortholog to define relative gains/losses in humans; and (2) against the human ancestral sequence to quantify divergence of ancestral-derived pairs. Differential activity was assessed in a pairwise manner with the testCoefficient() function (Wald test), and differential pairs were defined at a 5% FDR using the Benjamini-Hochberg procedure.

### ABC score

Regulatory activity for each TSS was calculated using the ABC score framework defined in (Fulco et al. 2019), substituting the estimated transcription rate □ from MPRAnalyze for activity and using (genomic distance)^-0.7^ as a proxy for contact frequency. CREs within 5 Mb of each TSS were considered for the calculation. While ABC values per CRE are typically normalized and interpreted as a relative contribution to a given TSS, we compared summed ABC scores between paralogous genes. Regulatory divergence was scored in a similar manner to expression divergence: a pseudocount of one order of magnitude below the smallest nonzero value was added to all ABC values, and the log_2_ ratio of scores was calculated for each derived-ancestral gene pair.

### ChIP-seq and ATAC-seq analyses

For calculation of ABC scores as published (using H3K27ac ChIP and ATAC data) and comparison with the MPRA, we aligned short reads (Supplemental Table S1) to GRCh38 with Bowtie v1.1.2 (Langmead et al. 2015), allowing for multiple mapping with the flags “-a -v2 -m 99”. CSEM was used to assign a probability to each alignment position on the basis of the local unique mapping rate (Zhang et al. 2020; Chung et al. 2014). Reads were then allocated to their most likely position by selecting the alignment with the highest posterior probability, or at random if all mappings were equally likely. Peaks were called on the ChIP data using input controls at a 5% FDR using MACS2 (Liu 2014). For ATAC and DHS data, the additional parameters “--nomodel --shift -100 --extsize 200 –broad” were added. Read counts per peak were determined with BEDTools coverage (Quinlan and Hall 2010).

### Luciferase reporter assays

1-kb fragments were synthesized by Azenta Life Sciences and cloned into the pE1B vector using the NEBuilder HiFI Assembly Master Mix (NEB #E2621). GM12878 and SH-SY5Y cells were cultured as described above. SH-SY5Y cells at 70-90% confluence were cotransfected with 50 ng pRL-TK and equimolar test construct using Lipofectamine™ 3000 (ThermoFisher L3000001). Cells were lysed 48 hours post-transfection and assayed with the Dual-Luciferase Reporter Assay System (Promega E1910) and the Tecan Spark plate reader according to the manufacturer’s instructions. GM12878 cells were split 48 and 24 hours prior to transfection and electroporated with 12.5 μg pRL-TK and equimolar test construct, brought to 100 μL with RPMI. For each transfection, 12.5 million cells were electroporated with the Neon system (ThermoFisher) using buffer E2 and the following program: 1200 V, 20 ms, 3 pulses. Cells were recovered in 4.3 mL of pre-warmed media (RPMI+15% FBS, no antibiotics). GFP controls were used to monitor transfection efficiency (∼15%). 24 hours post-transfection, cells were washed in PBS, plated at 500,000 cells per well, and lysed in an optical plate. Luminescence measurements were performed as above. The activity of each construct was quantified relative to the empty vector, and differential activity was relative to the human ancestral sequence. Significant differences were determined by ANOVA, using batch as a covariate, followed by a Tukey post-hoc test.

### DiMeLo-seq analyses

DiMeLo data for H3K4me3 and H3K27ac in GM12878 were obtained from GSE208125 (Maslan et al. 2024) as BAM files aligned to T2T CHM13 v2.0. Base modifications (5mC and 6mA) were thresholded at 155 (corresponding to probability of ∼60%) and the frequency of base modification per site was calculated using modkit pileup (Nanoporetech). For comparison with short-read ChIP data and MPRA tiles, coordinates were mapped to T2T CHM13 v2.0 using liftOver (UCSC Genome Browser Utilities). Then, the mean fraction of modified bases was calculated for each interval in all peak sets of interest using BEDTools map. The summed ABC score was calculated per HSD gene as above, using only the H3K27ac signal. Finally, differences in signal (mean fraction modified bases) for all assays (5mC, H3K27ac 6mA, and H3K4me3 6mA) were compared between chimpanzee-syntenic and nonsyntenic regions, also converted to CHM13 v2.0 coordinates.

## DATA ACCESS

Raw sequence data generated in this study have been submitted to the NCBI BioProject database (http://www.ncbi.nlm.nih.gov/bioproject) under accession number PRJEB98375. Code used for this project are available at Zenodo (https://doi.org/10.5281/zenodo.17240139) and as Supplemental Code.

## ACKNOWLEDGMENTS

We thank Drs. Fumitaka Inoue and Tal Ashuach for technical advice related to MPRA library construction and data processing, as well as Drs. Sierra Nishizaki and Daniela C. Soto for valuable discussion of the bioinformatic analyses performed in this study. We also acknowledge Dr. Anthony Antonellis for sharing the pE1B enhancer reporter Gateway plasmid, as well as Drs. Gerald Quon and Siobhan Brady for constructive feedback on the manuscript. Thank you to Dr. Nicolas Altemose for suggestions and code for analyzing the DiMeLo-seq data. Finally, we acknowledge The ENCODE Consortium, specifically the laboratories of Bradley Bernstein, J. Michael Cherry, Thomas Gingeras, Brenton Graveley, and John Stamatoyannopoulos for data used in this study. This work was supported by the UC Davis COVID-Impacted Research Funding Program, National Science Foundation (CAREER 2145885 to M.Y.D.), and National Human Genome Research Institute (F31HG011205 to C.S.) at the National Institutes of Health.

## AUTHOR CONTRIBUTIONS

C.J.S. and M.Y.D. conceived the project. C.J.S., G.K., and M.Y.D. designed the experiments and generated data. C.J.S., S.P.M., and M.Y.D. performed data analyses. All authors contributed to writing the manuscript.

