## Supplemental Materials for "Influence of *cis*-regulatory elements on expression divergence in human segmental duplications"

**This PDF file includes:**
Supplemental Notes S1–S4
Supplemental Figures S1–S21

### SUPPLEMENTAL NOTES

**Supplemental Note S1.** Assayed CREs identified from LCL and brain datasets were smaller than elements intersecting multiple tissue types (median size 700, 600, and 1400 base pairs, respectively) (Supplemental Fig. S5A). Grouping test oligos by dataset of origin, the largest number was unique to the fetal ATAC datasets (N=626), followed by LCL DNAse sites (N=591), and then regions identified by all datasets (N=582). We also stratified activity (alpha) scores by the number of supporting datasets and found a weak but significant relationship in the SH-SY5Y MPRA (additional 0.05 units of activity per brain dataset, $p<2.2\times10^{-16}$) (Supplemental Fig. S5B). In GM12878, there was an even weaker trend (additional 0.004 units of activity per LCL dataset, $p=0.07$) (Supplemental Fig. S5B). Thus, an increasing number of datasets does not necessarily converge on stronger regulatory elements.

We also considered differences in activity arising from inexact alignment lengths between ancestral and paralogous or orthologous regions, which resulted in some alignments being trimmed or padded within 5% tolerance. In both cell types, the activity of exact-length sequences was slightly lower (0.98-fold difference in SH-SY5Y, Wilcoxon rank sum test $p=1.2\times10^{-5}$; 0.97-fold difference in GM12878, $p<2\times10^{-16}$). There was nominally a quantitative effect of the difference in number of bases in GM12878, but the correlation was very low ($r=-0.04$, $p=5\times10^{-4}$). Meanwhile, there were no differences in the magnitude of differential activity between exact and trimmed/padded sequences, and a quantitative effect of a similar scale (Supplemental Fig. S6). We do not expect that trimming or padding had a systematic effect on sequence activity, as unmodified sequences had a larger range and comprised the majority of the oligos tested.

**Supplemental Note S2.** To examine the discrepancy between expression datasets in the correlation with ABC score ratios, we downsampled the short-read SH-SY5Y data to match the long-read Kinnex read count. After reducing the short-read SH-SY5Y data to 32% and only considering replicates individually, there is still a significant correlation of expression divergence with ABC score differences ($p<0.04$, $r>0.43$ for all replicates; Supplemental Fig. S15). Differences are thus unlikely to be a result of coverage, but other biases inherent to short- versus long-read sequencing. For example, in Illumina datasets, one transcript generates multiple fragments, while in Kinnex one transcript yields one count. Full-length transcripts must also be completely reverse-transcribed and amplified. Indeed, long-read expression datasets had a larger fraction of undetected genes than short-read datasets, which persisted after downsampling (Table S2). Finally, the calculation of expression divergence as a ratio of two expression values (added to a pseudocount and $\log_2$-transformed) is sensitive to low or missing value, and this metric had a poorer correlation across gene pairs than individual genes (Supplemental Fig. S1C–D). For example, NAIPP1, NAIPP2, and OCLNP1 were lowly expressed in short-read data and not detected with long read, resulting in much lower expression divergence scores.

**Supplemental Note S3.** Using histone modification (H3K4me3, H3K27ac) data from directed methylation with long-read sequencing (DiMeLo-seq) assays (Maslan et al. 2024) in GM12878, we found an expected correlation between HSD gene expression and histone H3 active marks; compared to short-read ChIP this relationship was stronger and also tracked with the expression differences between paralogous pairs, when coupled with the long-read expression quantification (Supplemental Fig. S20A). We also found that CpG methylation at HSD promoters inversely correlated with expression overall, but not for distinguishing ancestral-derived gene pairs (Supplemental Fig. S20A). While promoter activity itself did not show this pattern in the MPRA, the chromatin state captured at promoters does not cause higher transcription *per se*, instead reflecting a downstream output of total CRE activity. Instead, this showcases the increased accuracy on the long-read DiMeLo assays relative to ChIP and supports the expression analyses.

Globally, H3K4me3 and 5mC signals were higher in chimpanzee-syntenic (likely ancestral) than non-syntenic (likely derived) regions, (respective 1.08- and 1.19-fold difference of medians), consistent with greater promoter activity and transcription-associated gene body methylation (Supplemental Fig. S20B). Interestingly, 5mC was not different when considering only promoters, counter to previous reports that CpG methylation tracked with expression in a few duplicated gene families (Vollger et al. 2022). Taken together, these results are in accord with observed differences in RNA levels.

We again calculated an ABC score per gene using DiMeLo signal at the same peaks defined with ChIP-seq. In agreement with the ChIP-based analysis in SH-SY5Y, ABC score ratios from DiMeLo in GM12878 correlated with expression divergence (short-read RNA, Figure 5). This result held when subsetting to only peaks in unique regions, but no correlation was found using HSD peaks only. This again suggests a larger contribution of adjacent, non-duplicated sequence to expression differences among HSD paralogs.

**Supplemental Note S4.** Due to the quality and completeness of the telomere-to-telomere (T2T) CHM13 assembly (hs1), we tabulated differences from our test oligos designed to GRCh38 (hg38). Our focal duplicated regions were resolved to comparable accuracy (Dennis et al. 2017) and accordingly represent an alternative haplotype. We performed a liftOver from hg38 to T2T-CHM13 and found that 3,891/5,716 (68%) of tiles were identical in both assemblies, (Supplemental Fig. S21A). The remaining tiles failed to lift over, were not in hg38 (*DUSP22* and *GPRIN2* contigs), or had a mix of identical and non-identical matches (i.e. identical paralogous positions in hg38 that were not all identical in T2T-CHM13). Excluding these, 74% were identical. Similarly, we compared our 2,444 chimpanzee sequence derived from panTro6 to the highly contiguous mPanTro3, haplotype 1 (Yoo et al. 2025, HPRCy1 chain). Of these, 1,467 (61%) lifted over with exact sequence matches (Supplemental Fig. S21A). For reference, a single nucleotide divergence rate of 0.1% or 0.2% corresponds to a predicted identical fraction of 81.9% or 72.6%, respectively (Poisson expectation). There was no skew for promoters versus distal elements in any of the categories in either species (Fisher's Exact Test).

To consider how these differences might affect our analysis, we compared measured MPRA activity between sequences that were identical in both genomes or not. For human tiles in both cell types, the non-identical sequences had nominally higher activity (Wilcoxon rank-sum test, $p<0.02$), but the magnitude was small (median fold difference ~1.01) (Supplemental Fig. S21B). The pattern was opposite for chimpanzee sequences ($p<0.02$, median fold difference >0.98). Differential activity, scored as the absolute value of the $\log_2$-fold difference of sequence pairs, was stronger in identical sequences in SH-SY5Y ($p=8\times10^{-3}$, median fold difference 1.23, Supplemental Fig. S21B), though this effect disappeared when considering only differentially active (5% FDR) pairs. In the chimpanzee comparison, differential activity was stronger in non-identical sequences in GM12878 ($p=0.02$, median fold difference 0.92). While allelic variation almost certainly will result in differences in regulatory activity, we do not expect these to affect any of the patterns reported in this study. Cross-genome comparisons of sequence content are annotated in Supplemental Table S6.

#### SUPPLEMENTAL FIGURES

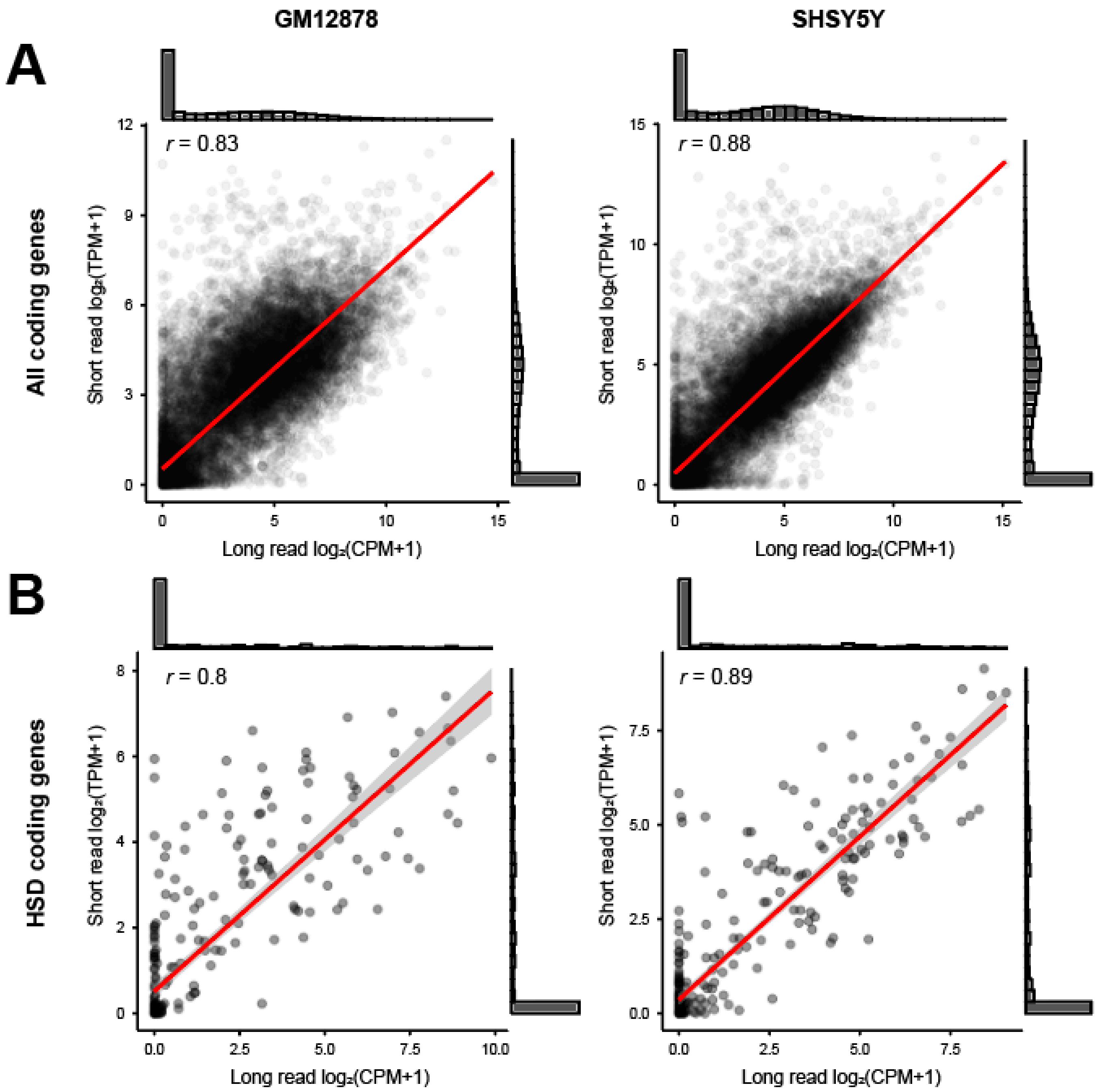

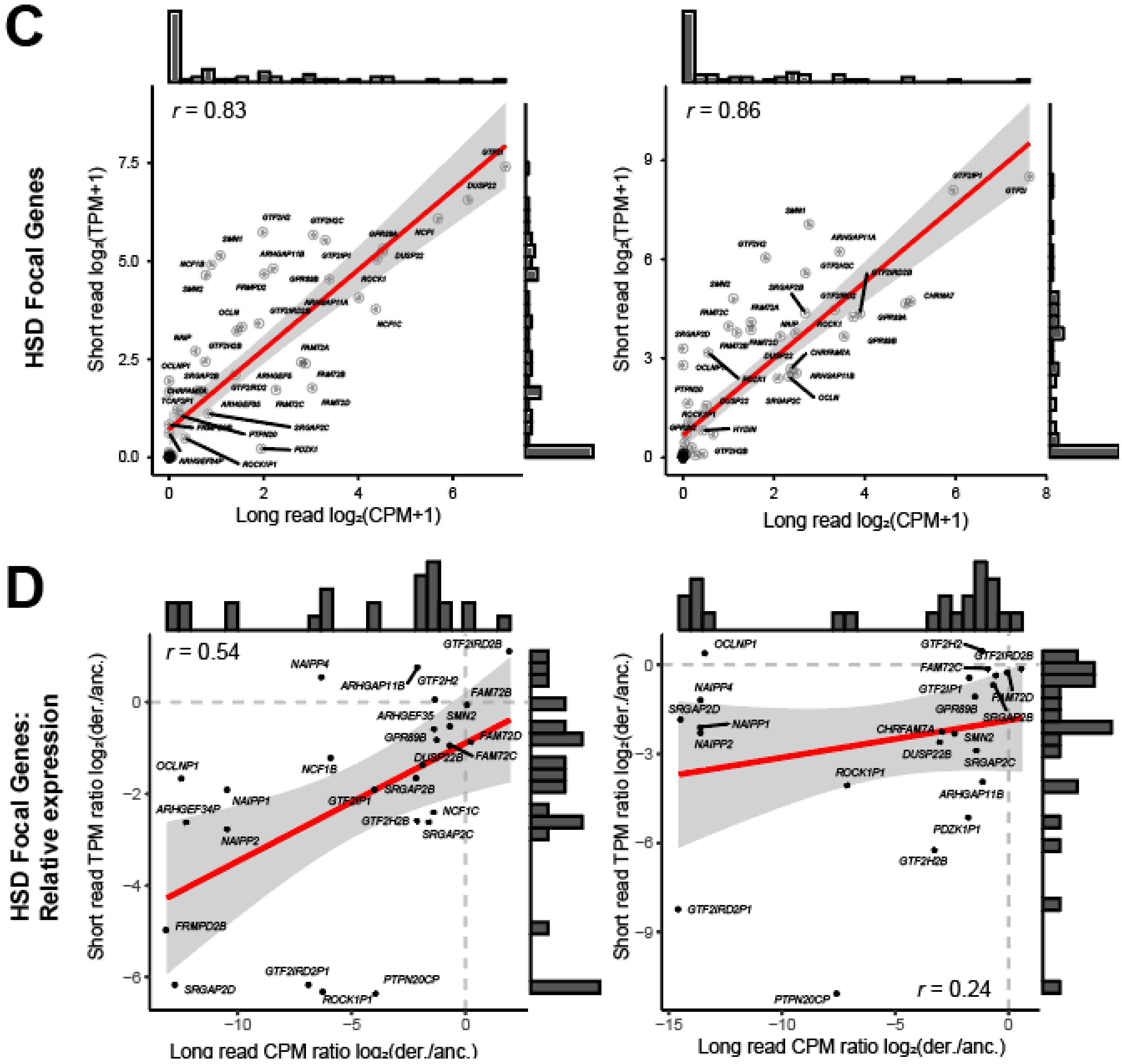


**Supplemental Fig. S1. Short- and long-read expression data are correlated.** Comparison of $\log_2$-transformed expression data from short-read RNA sequencing (Illumina) and long-read RNA sequencing (PacBio Kinnex) for polyadenylated genes (non-histone coding genes) detectable by Kinnex: **(A)** all genes, **(B)** HSD genes. Focal HSD genes for this study, which are all polyadenylated, are shown in **(C)**. **(D)** Expression divergence for ancestral-derived pairs of HSD focal genes, calculated as the $\log_2$ ratio of derived (der.) to ancestral (anc.) expression, with a pseudocount (one order of magnitude smaller than the lowest nonzero expression) added to each. Only pairs expressed above 1 TPM or 1 CPM are shown. The Pearson's correlation *r* is printed on each plot.

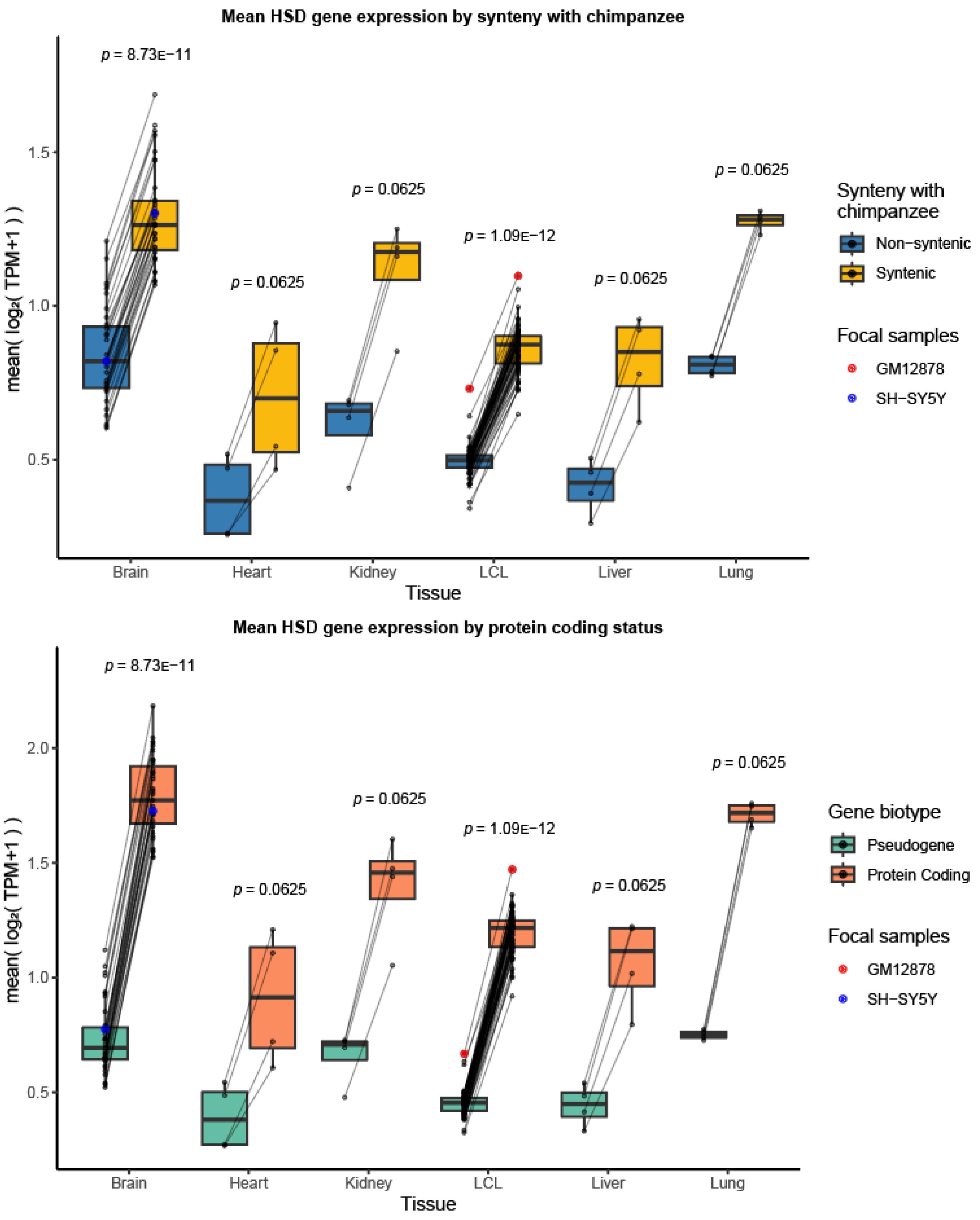


**Supplemental Fig. S2. HSD genes are differentially expressed across tissues.** Mean expression for HSD gene sets in the cell lines used in this work (SH-SY5Y and GM12878), additional lymphoblastoid cell lines, and primary brain, heart, kidney, liver, and lung. Gene sets are divided by **(A)** ancestral or derived status as defined by chimpanzee synteny and **(B)** gene biotype (protein coding or pseudogene). Each sample is represented by a pair of points. *P*-values shown were determined by a paired Wilcoxon signed-rank text with Benjamini–Hochberg (BH) adjustment.

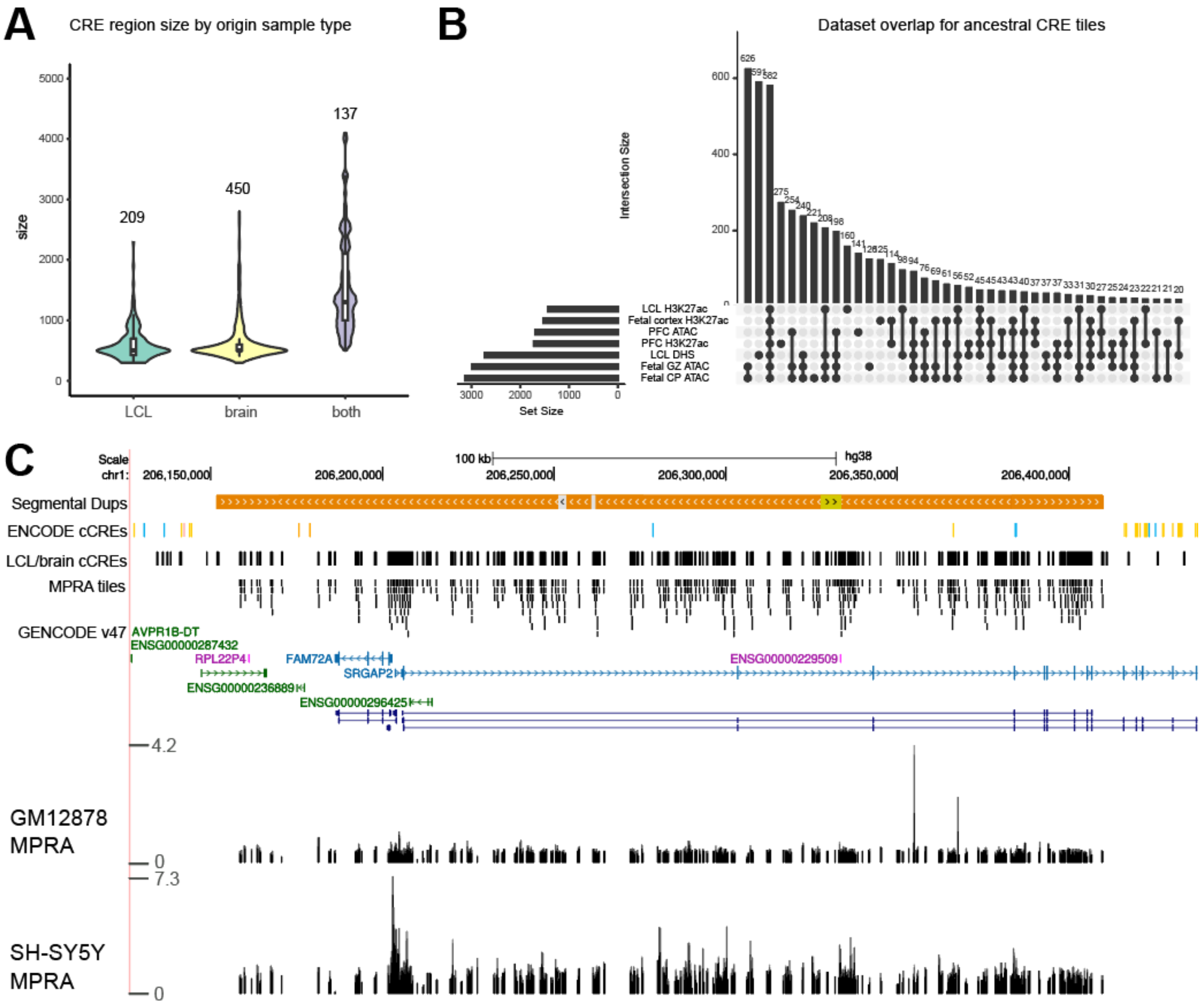


**Supplemental Fig. S3. MPRA design. (A)** Distribution of merged CRE region sizes used to design MPRA tiles, separated by overlap with original dataset tissue types. The size of each set is printed over each violin plot. **(B)** Intersections of 200mer tiles with original datasets. **(C)** Example locus showing *FAM72A* and *SRGAP2* Chromosome 1. Segmental duplications (Segmental Dups) of >99% nucleotide identity are shown as orange bars. ENCODE candidate *cis*-regulatory elements (cCREs) are shown as published: promoters (red), proximal enhancers (orange), distal enhancers (yellow), CTCF only (blue). Duplicated cCREs as developed by this study are shown in black, in addition to the exact tiles measured in MPRA. Beneath the gene annotation (GENCODE v47), sequence activity as determined by MPRA is shown for each cell type as the alpha value for each 200mer tile assayed.

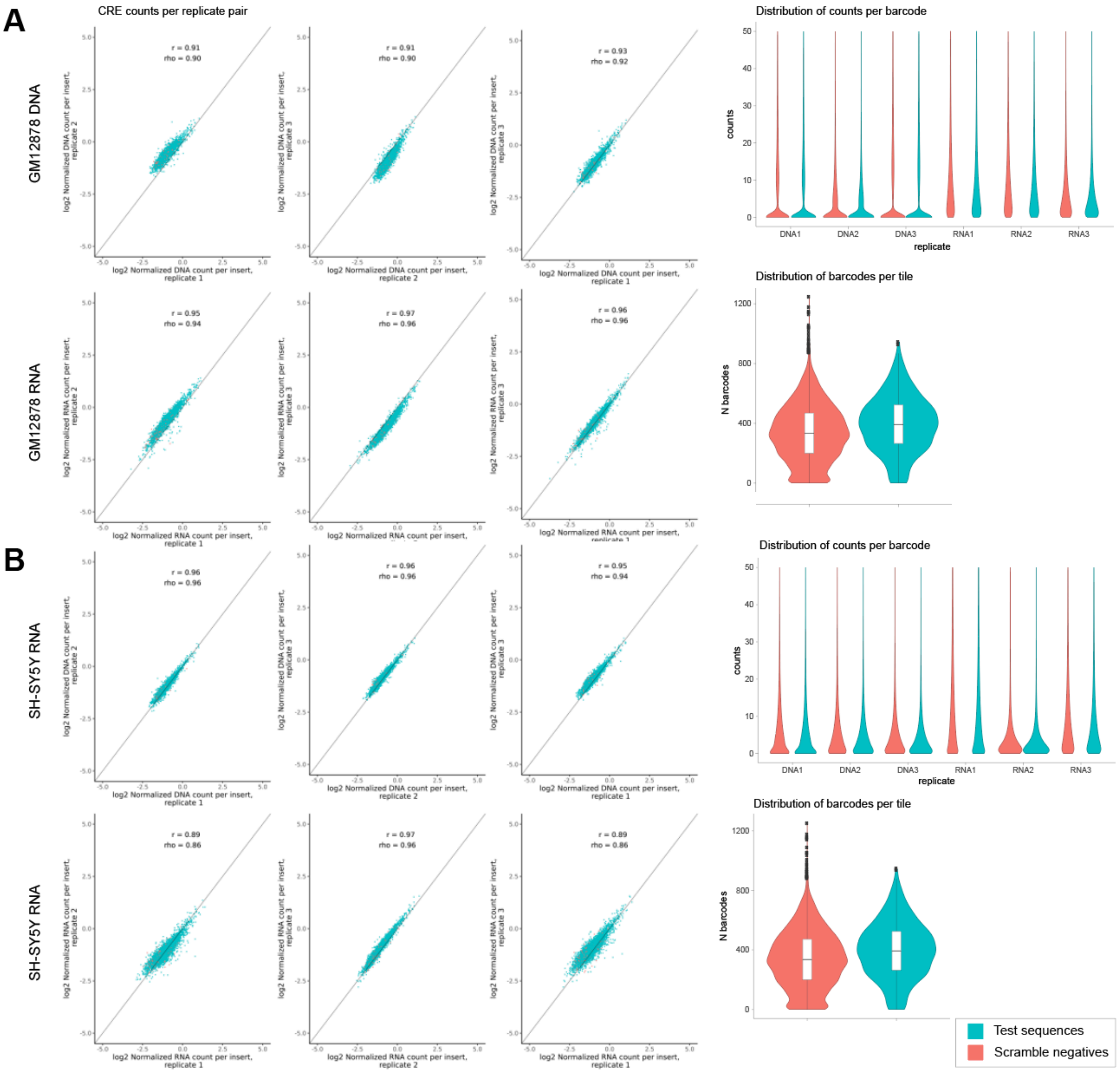


**Supplemental Fig. S4. MPRA is reproducible.** MPRA summary statistics for **(A)** GM12878 and **(B)** SH-SY5Y. Test sequences designed to HSDs are shown in turquoise, while scramble negative controls are shown in salmon. Left: Comparison of log-normalized counts for all pairs of replicates, for DNA and cDNA (RNA) libraries in each cell type. Each scatter plot is annotated with Spearman and Pearson's correlation coefficients. Top right: Distributions of sequencing counts per barcode for all replicates. Bottom right: Distribution of barcodes counted per MPRA insert (tile).

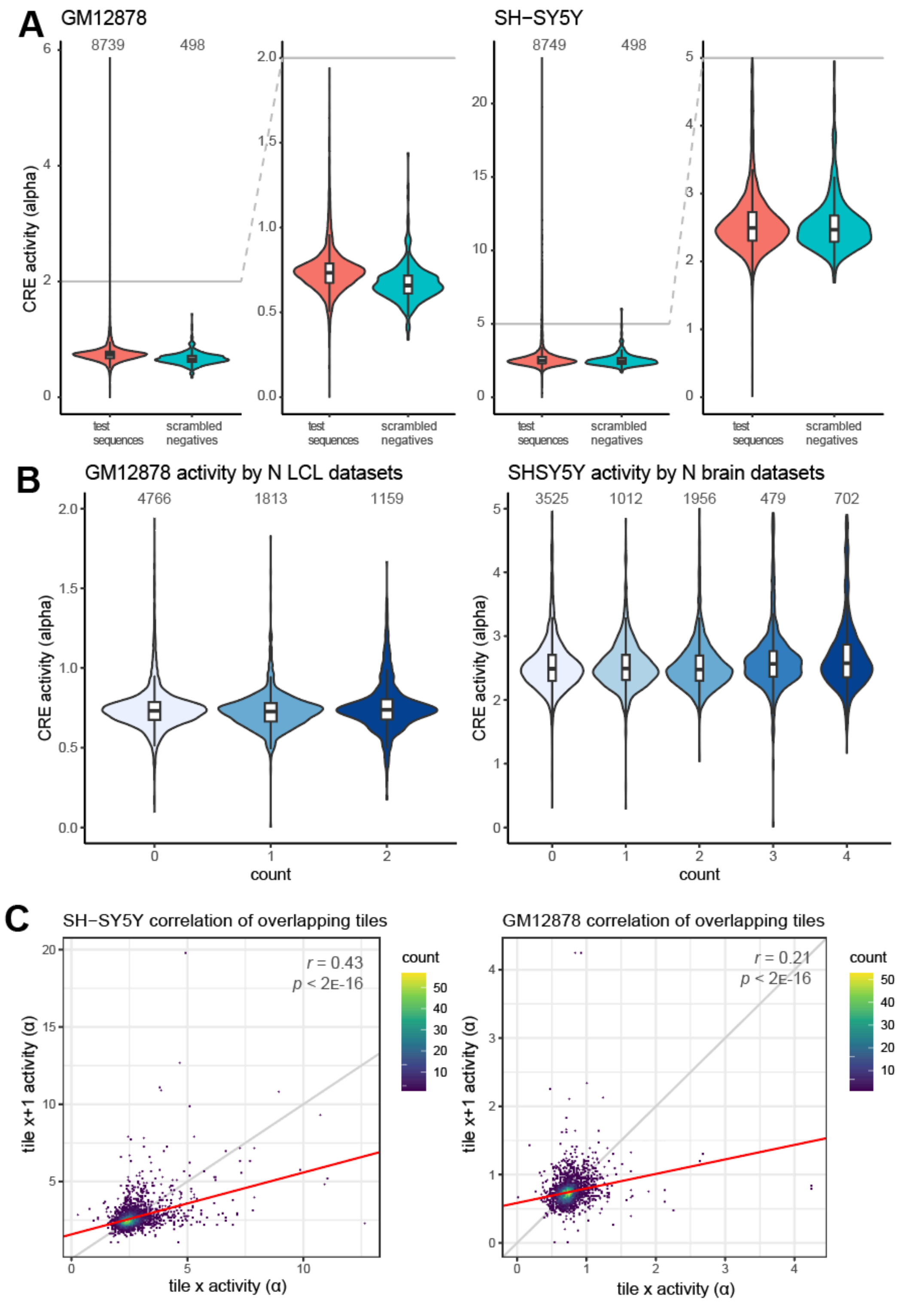


**Supplemental Fig. S5. MPRA activity measurements.** MPRA activity measurements from experiments performed in GM12878 (left) and SH-SY5Y (right) **(A)** Distribution of alpha values, separated for test HSD sequences and negative controls. The number of CREs per set is printed over each violin plot. Each plot is reproduced on the right with a cropped y-axis. **(B)** Distribution of alpha values separated by the number of supporting datasets in each cell type. There was a weak linear association in SH-SY5Y (additional 0.05 units of activity per brain dataset, $p<2.2\times10^{-16}$). **(C)** Scatter plots of CRE activity for all pairs of overlapping tiles in each cell type. The line of best fit is shown in red, while the identity y=x is printed in gray. The Pearson's correlation coefficient and *p*-value for linear association are displayed in the top right corner of each plot.

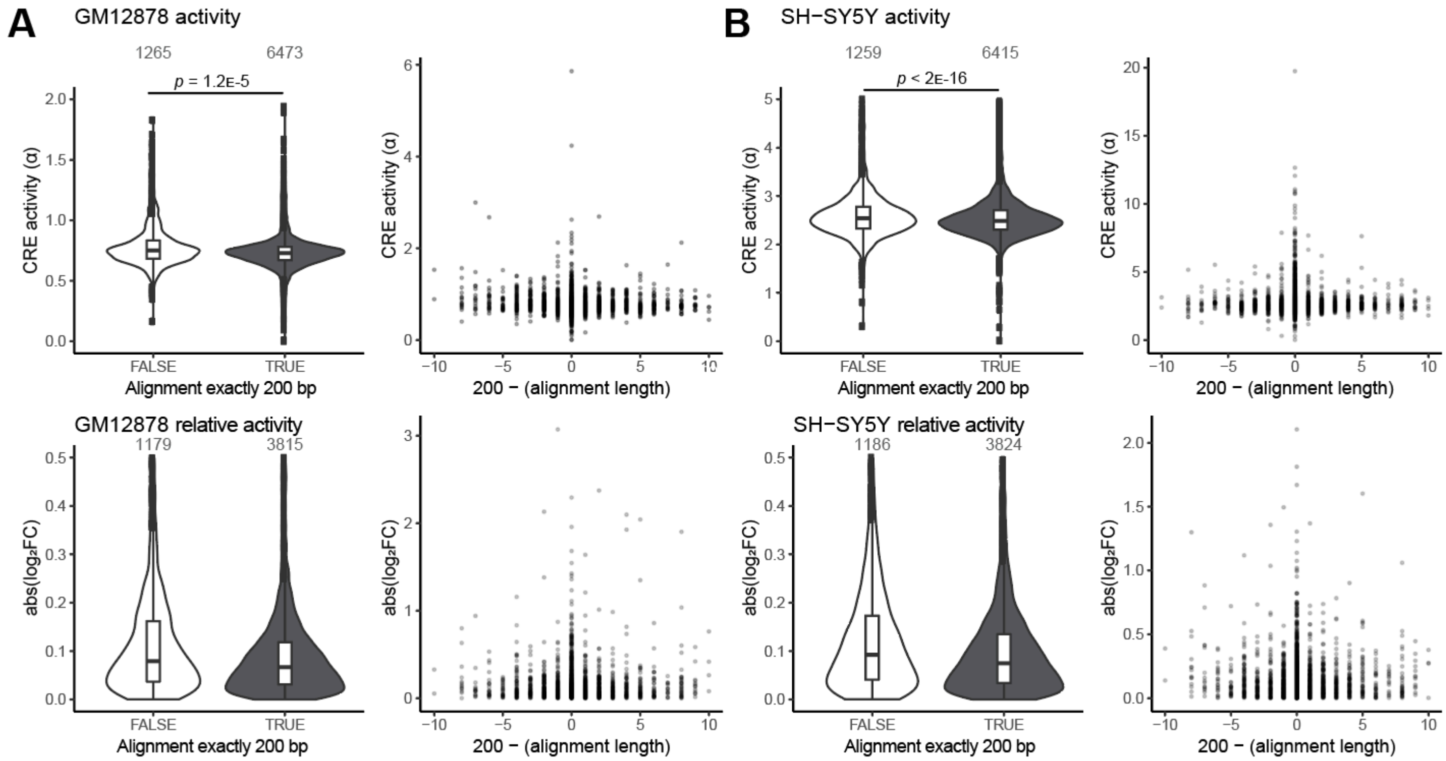


**Supplemental Fig. S6. Comparison of MPRA sequences by alignment length prior to trimming or padding.** All tiles designed in ancestral regions had a size of 200 bp, while aligned regions in derived paralogs or chimpanzee orthologs were allowed a 5% tolerance. MPRA in GM12878 is shown in **(A)** and SH-SY5Y is shown in **(B)**. Left: Activity of all sequences and relative activity of derived-ancestral pairs, separated by the size of the original alignment length. The number of sequences in each group is printed above each violin plot. Right: Activity and relative activity plotted by difference in original alignment size.

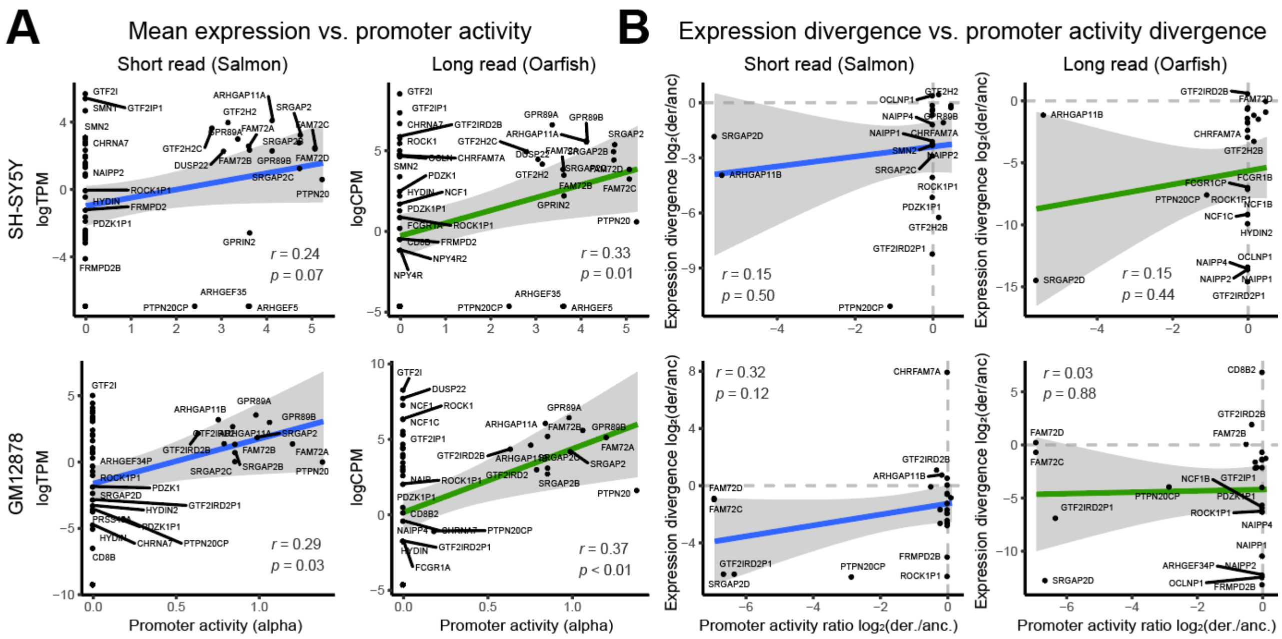


**Supplemental Fig. S7. MPRA at promoters does not capture all regulatory information. (A)** Pearson's correlation of log-transformed gene expression (plus a pseudocount one order of magnitude below the smallest nonzero value, logTPM or logCPM) and promoter activity (mean alpha value within 1 kb of the TSS). Each cell type and gene expression dataset are plotted separately. **(B)** Pearson's correlation of relative expression for each derived/ancestral (der./anc.) gene pair and relative promoter activity. Only expressed gene pairs are included, and ratios are $\log_2$-transformed. The 95% confidence interval for the regression is shaded in gray.

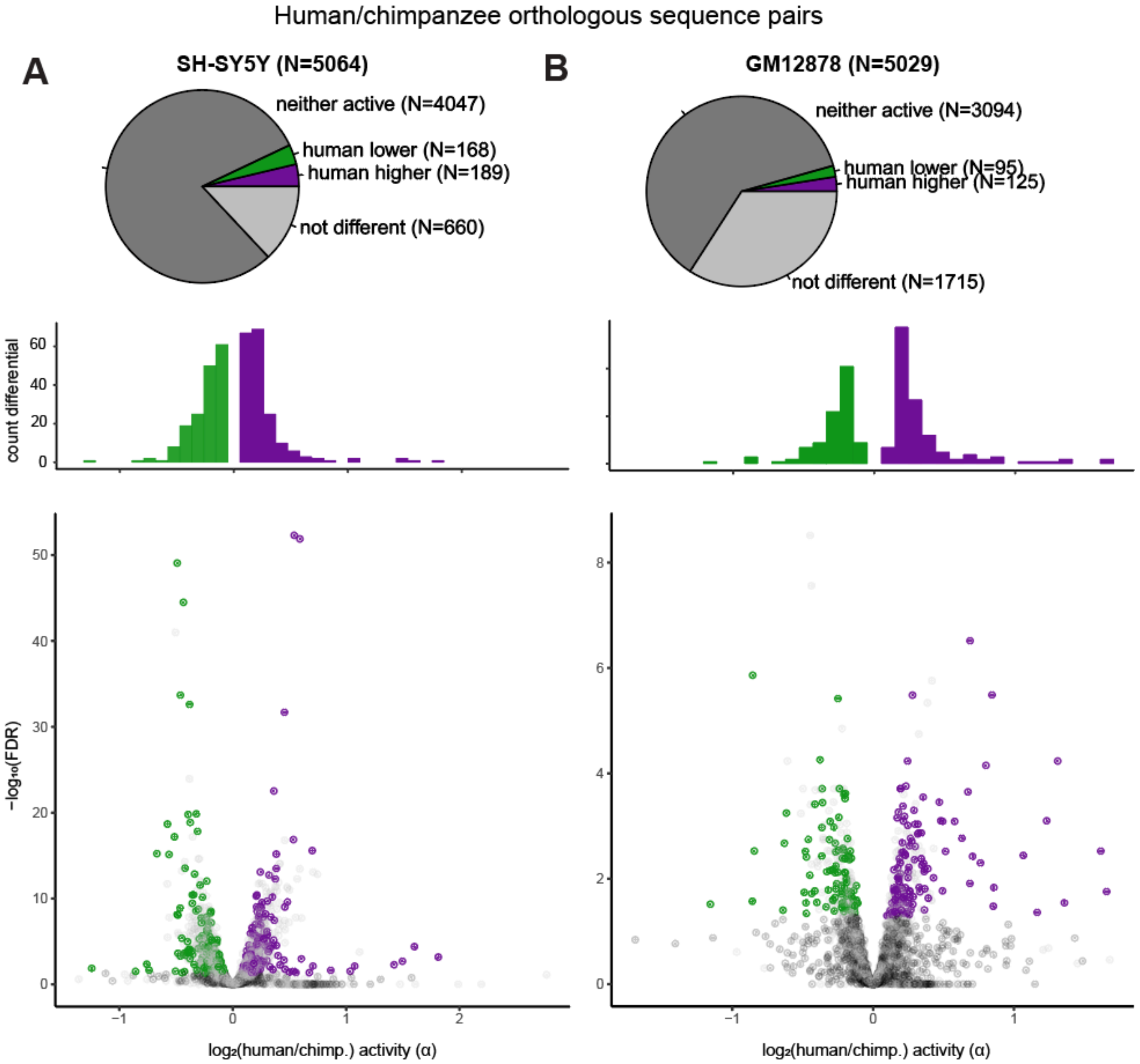


**Supplemental Fig. S8. Comparison of sequence activity between species. (A)** Sequence activity in SH-SY5Y, relative to the chimpanzee (chimp.) ortholog. Pie charts tabulate human-chimpanzee sequence pairs as follows: green (human lower), purple (human higher), gray (not differential), black (neither sequence active). Volcano plots depict differential activity between human-chimpanzee orthologous sequence pairs, colored by the same scheme. The marginal histograms depict the distribution of log-fold differences for differentially active sequences. **(B)** Differential activity in GM12878.

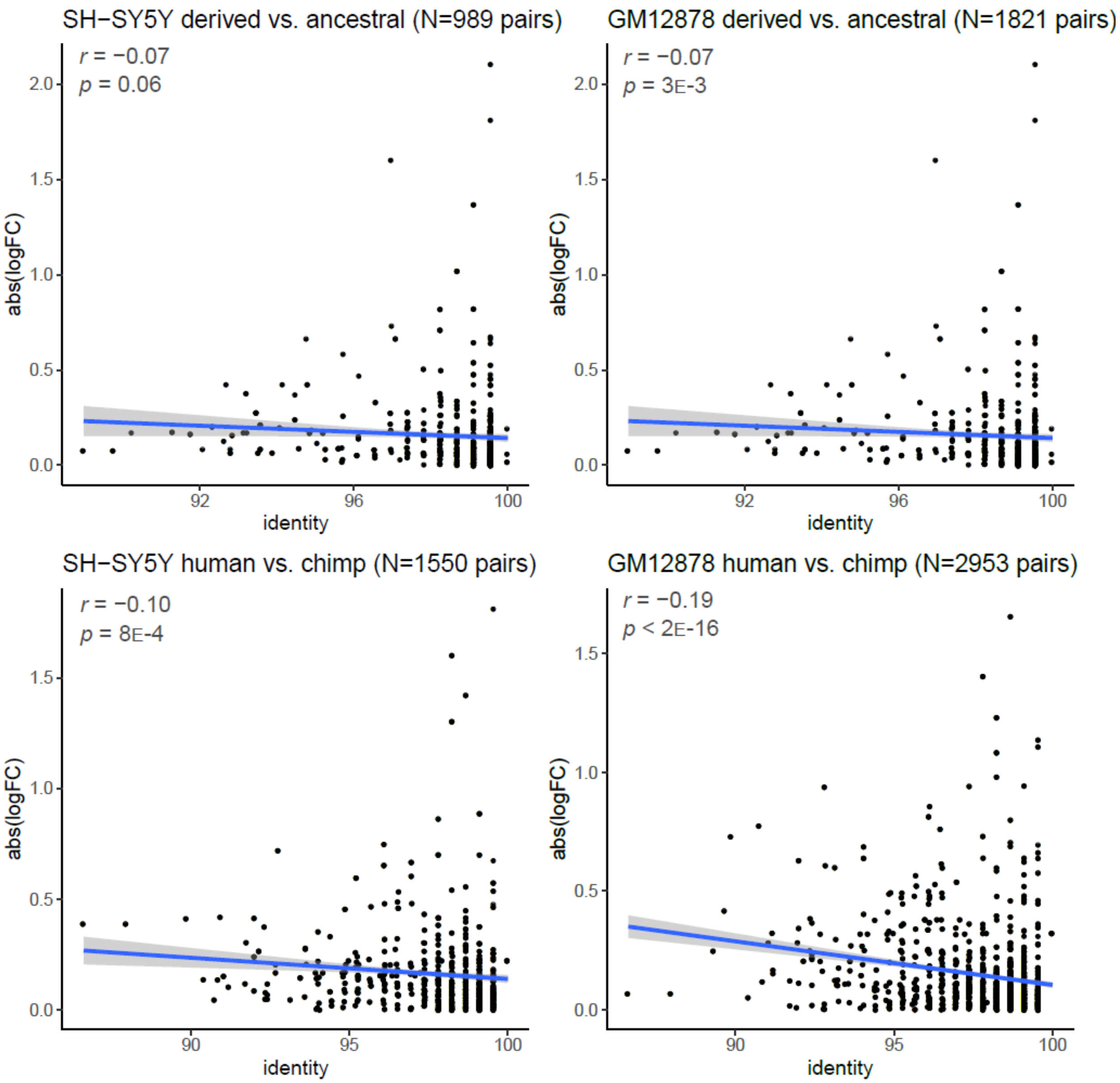


**Supplemental Fig. S9. Comparison of regulatory divergence and sequence identity.** For all sets of active sequence pairs (human derived/ancestral or human/chimpanzee) in each cell type, the absolute value of the $\log_2$-fold difference (logFC) is plotted against sequence identity, the percentage of identical nucleotides in each pair. The Pearson's correlation coefficient and *p*-value for the correlation are printed in the upper left of each plot, and the 95% confidence interval for the regression is shaded in gray.

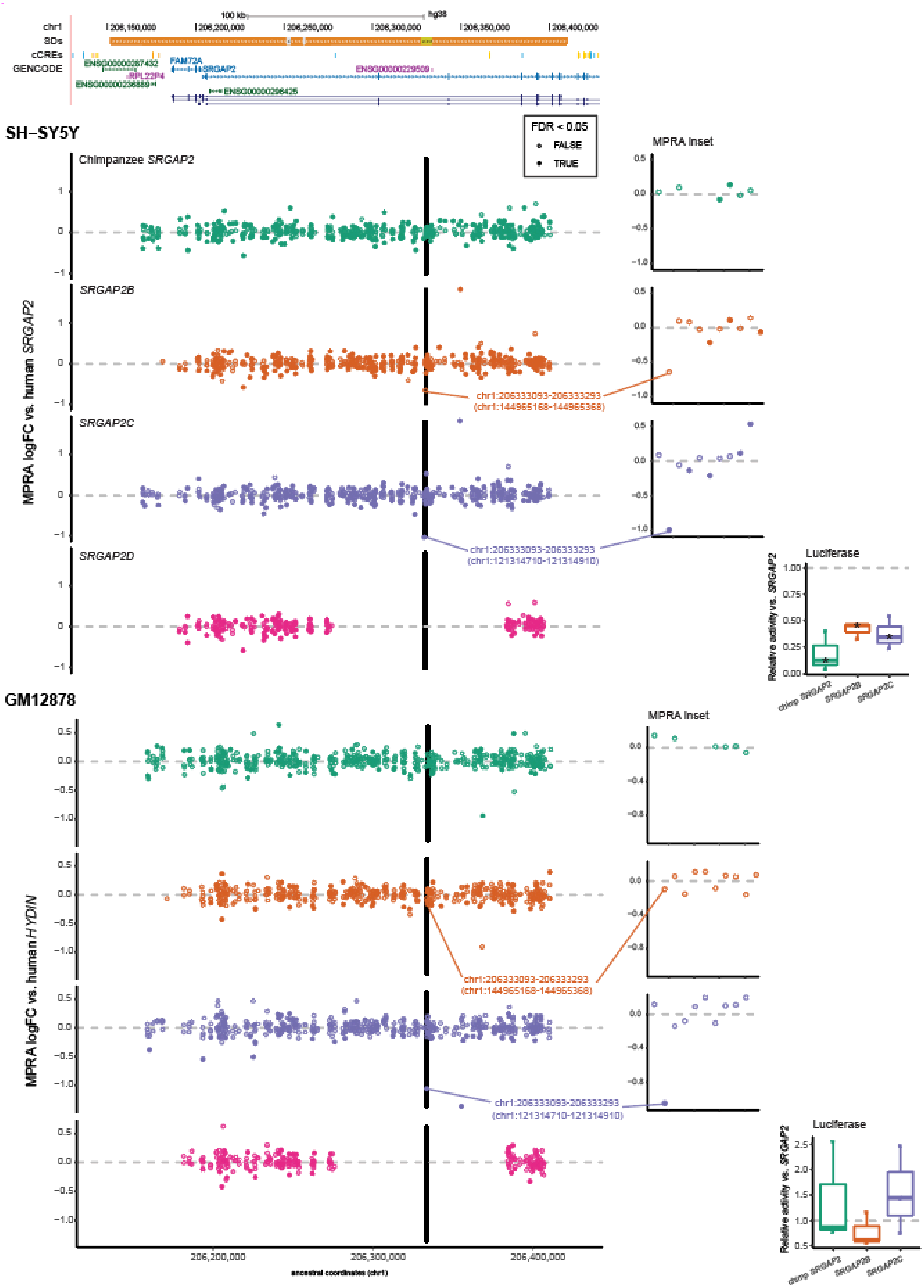

100 kb
hg38
chr1
SDs
cCREs
GENCODE
206,150,000
206,200,000
206,250,000
206,300,000
206,350,000
206,400,000
ENSG00000287432
FAM72A
SRGAP2
ENSG00000229509
RPL22P4
ENSG00000236889
ENSG00000298425
SH–SY5Y
FDR < 0.05
FALSE
TRUE
Chimpanzee SRGAP2
SRGAP2B
SRGAP2C
SRGAP2D
MPRA logFC vs. human SRGAP2
MPRA inset
chr1:206333093-206333293
(chr1:144965168-144965368)
chr1:206333093-206333293
(chr1:121314710-121314910)
Luciferase
Relative activity vs. SRGAP2
chimp SRGAP2
SRGAP2B
SRGAP2C
GM12878
MPRA logFC vs. human HYDIN
MPRA inset
chr1:206333093-206333293
(chr1:144965168-144965368)
chr1:206333093-206333293
(chr1:121314710-121314910)
Luciferase
Relative activity vs. SRGAP2
chimp SRGAP2
SRGAP2B
SRGAP2C
206,200,000
206,300,000
206,400,000
ancestral coordinates (chr1)

**Supplemental Fig. S10. Paralog-specific regulation in the *FAM72/SRGAP2* locus (Chr 1q32.1).** A 260-kb region (Chr 1:206,129,599-206,428,595; GRCh38) is shown. Segmental duplications (SDs) are colored orange for >99% sequence identity, and gray for <98% identity. ENCODE candidate *cis*-regulatory elements (cCREs) are shown as published: promoters (red), proximal enhancers (orange), distal enhancers (yellow), CTCF only (blue). MPRA data are visualized as activity relative to the human ancestral locus depicted in the browser coordinates. Each point represents a 200mer tile, and tiles are colored by which homolog they correspond to (teal: chimpanzee; orange: derived *SRGAP2B*; purple: derived *SRGAP2C*; pink: derived *SRGAP2D*) and filled in if they scored as differentially active. One tile differentially active in *SRGAP2C* is labeled, with the ancestral coordinates above and derived coordinates corresponding to the tested sequence underneath in parentheses. A 1-kb region is highlighted in the insets; the panel to the right shows a zoomed in plot of the MPRA measurements per tile, and the box plot shows a relative luciferase activity of the entire sequence, as assayed in the corresponding cell type. Asterisks on the box plots mark significantly different activity with respect to the human ancestral sequence. See also Table S13.

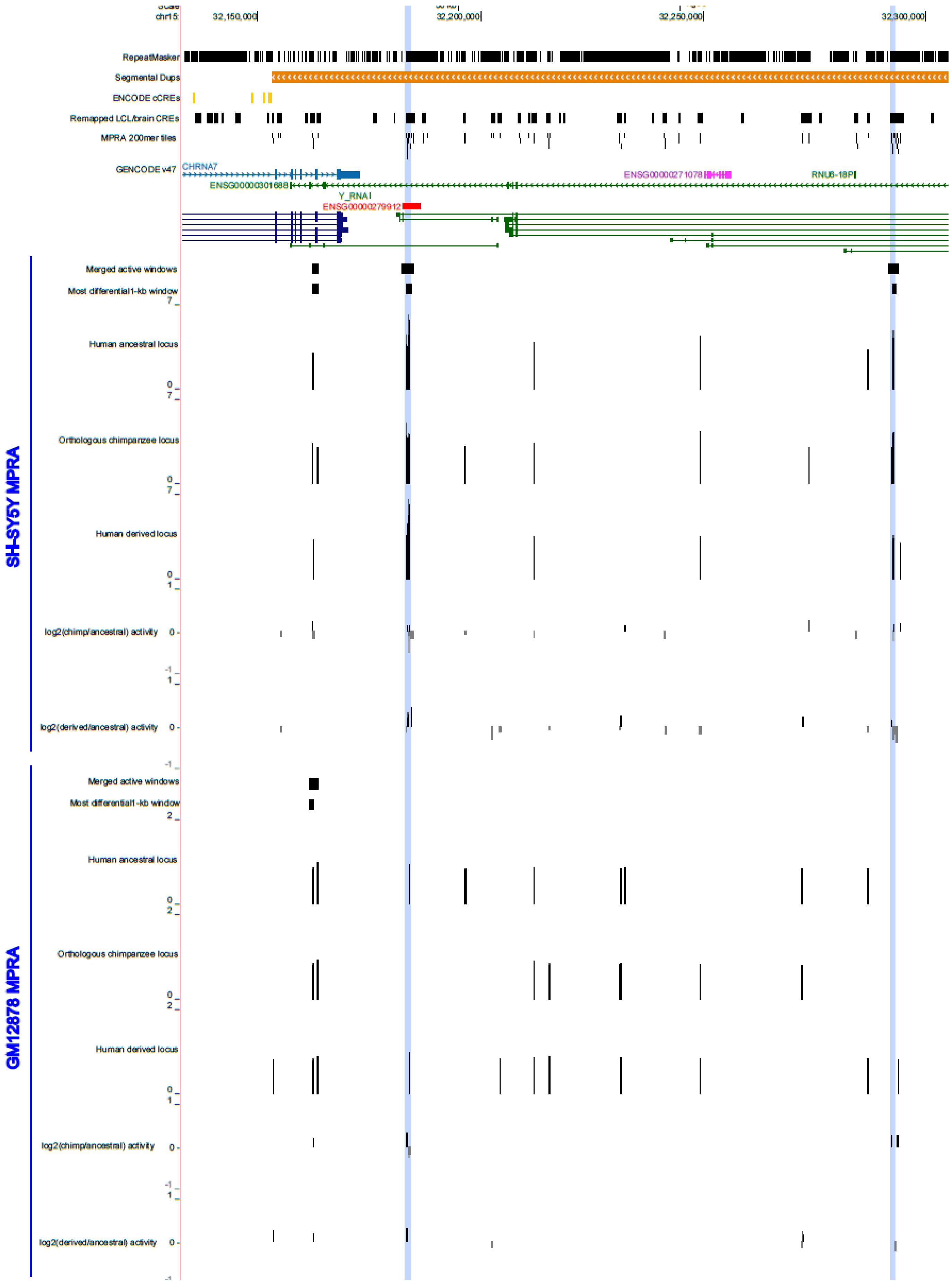
chr15:
32,150,000
32,200,000
32,250,000
32,300,000
RepeatMasker
Segmental Dups
ENCODE cCREs
Remapped LCL/brain CREs
MPRA 200mer tiles
GENCODE v47
CHRNA7
ENSG00000301688
Y_RNA
ENSG00000279912
ENSG00000271078
RNU6-18P
SH-SY5Y MPRA
Merged active windows
Most differential 1-kb window
Human ancestral locus
Orthologous chimpanzee locus
Human derived locus
log2(chimp/ancestral) activity
log2(derived/ancestral) activity
GM12878 MPRA
Merged active windows
Most differential 1-kb window
Human ancestral locus
Orthologous chimpanzee locus
Human derived locus
log2(chimp/ancestral) activity
log2(derived/ancestral) activity

**Supplemental Fig. S11. Example region (Chr 15q13.3) selected for cloning and luciferase assay**. Tracks from top to bottom: RepeatMasker annotation of transposable elements and simple repeats; segmental duplications, with duplicated sequence >99% identity shown in orange; ENCODE candidate *cis*-regulatory elements; merged human LCL and brain CRE used to design MPRA; tested MPRA 200mer tiles; GENCODE v47 annotation; for each MPRA, active merged regions, generated from top-scoring 1-kb windows; the single 1-kb window for each sequence overlapping the most strongly differential CRE between paralogs; human ancestral paralog (downstream of *CHRNA7*) activity; chimpanzee ortholog (*CHRNA7*) activity; human derived paralog (*CHRFAM7A*) activity; differential activity relative to chimpanzee *CHRNA7* ($log_2$-fold difference); differential activity relatives to human *CHRNA7* ($log_2$-fold difference). 1-kb sequences were nominated for having strong activity across tiles and selected (purple highlight) for containing multiple differentially active tiles.

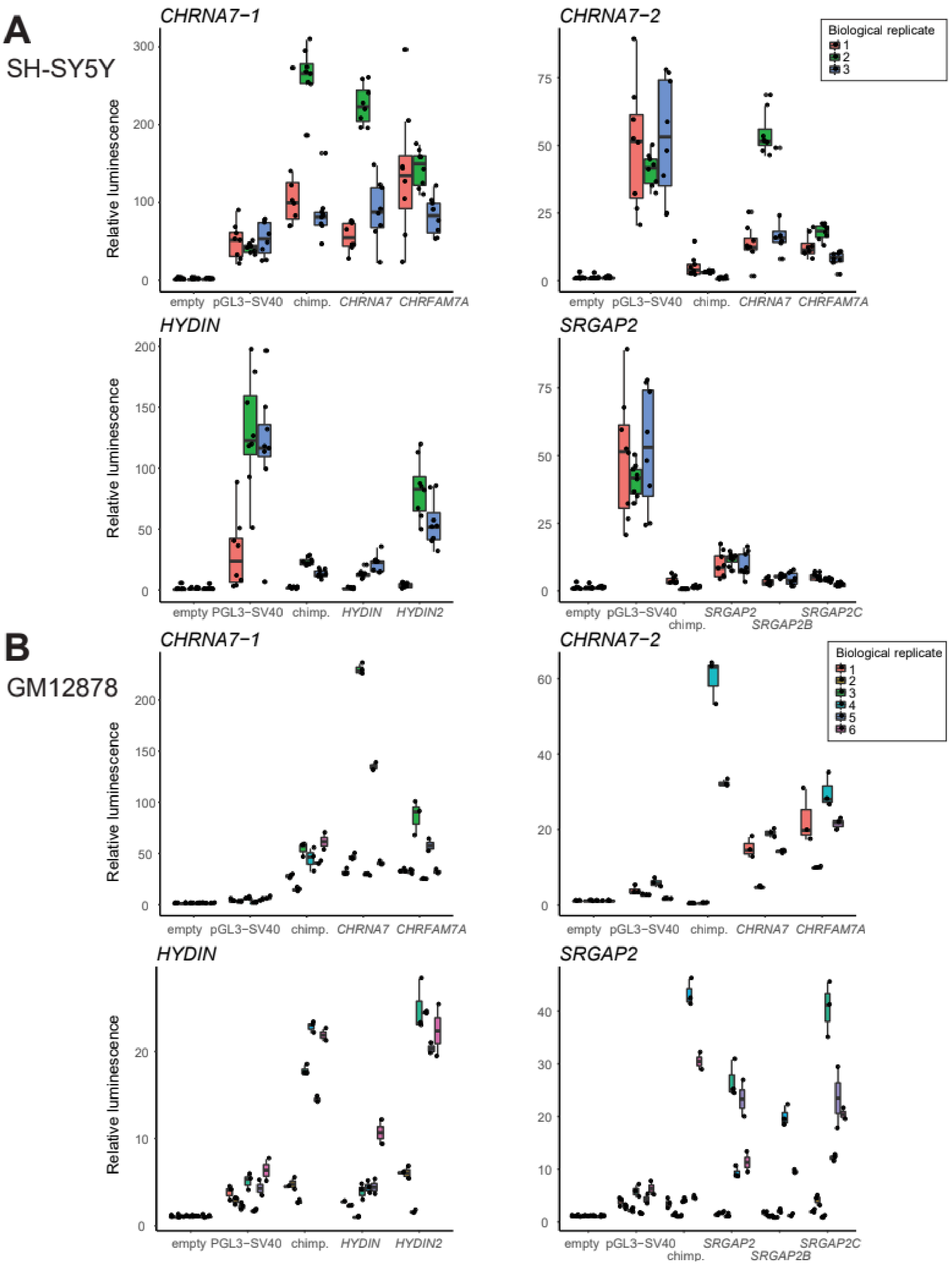


**Supplemental Fig. S12. Luciferase assay data for MPRA-nominated 1-kb regions.** Luminescence is plotted relative to the mean empty control value in each batch. Human paralogs are shown by gene name, and the chimpanzee ortholog is abbreviated as "chimp." The pGL3-SV40 positive control is shown for reference. **(A)** SH-SY5Y, **(B)** GM12878.

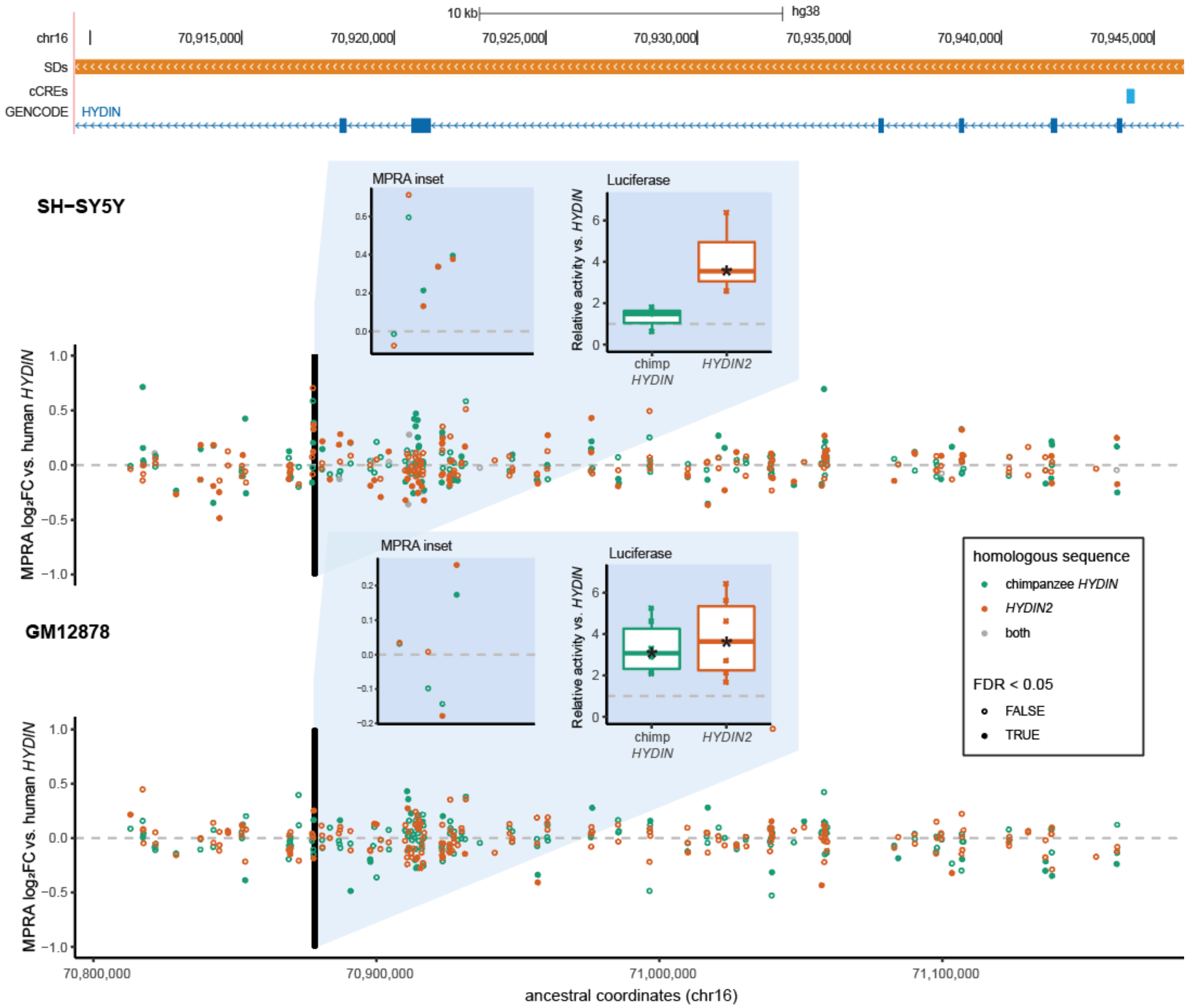


**Supplemental Fig. S13. Paralog-specific regulation in the *HYDIN* locus (Chr 16q22.2).** A 429-kb region (Chr 16:70,811,384-71,168,670; GRCh38) is shown. Segmental duplications (SDs) are colored orange for >99% sequence identity, and gray for <98% identity. ENCODE candidate *cis*-regulatory elements (cCREs) are shown as published: promoters (red), proximal enhancers (orange), distal enhancers (yellow), CTCF only (blue). MPRA data are visualized as activity relative to the human ancestral locus depicted in the browser coordinates. Each point represents a 200mer tile, and tiles are color by which homolog they correspond to (teal: chimpanzee; orange: derived *HYDIN2*; gray: identical in both) and filled in if they scored as differentially active. A 1-kb region is highlighted in the insets; the left panel shows a zoomed in plot of the MPRA measurements per tile, and the right shows a relative luciferase activity of the entire sequence, as assayed in the corresponding cell type. Asterisks on the box plots mark significantly different activity with respect to the human ancestral sequence. See also Table S13.

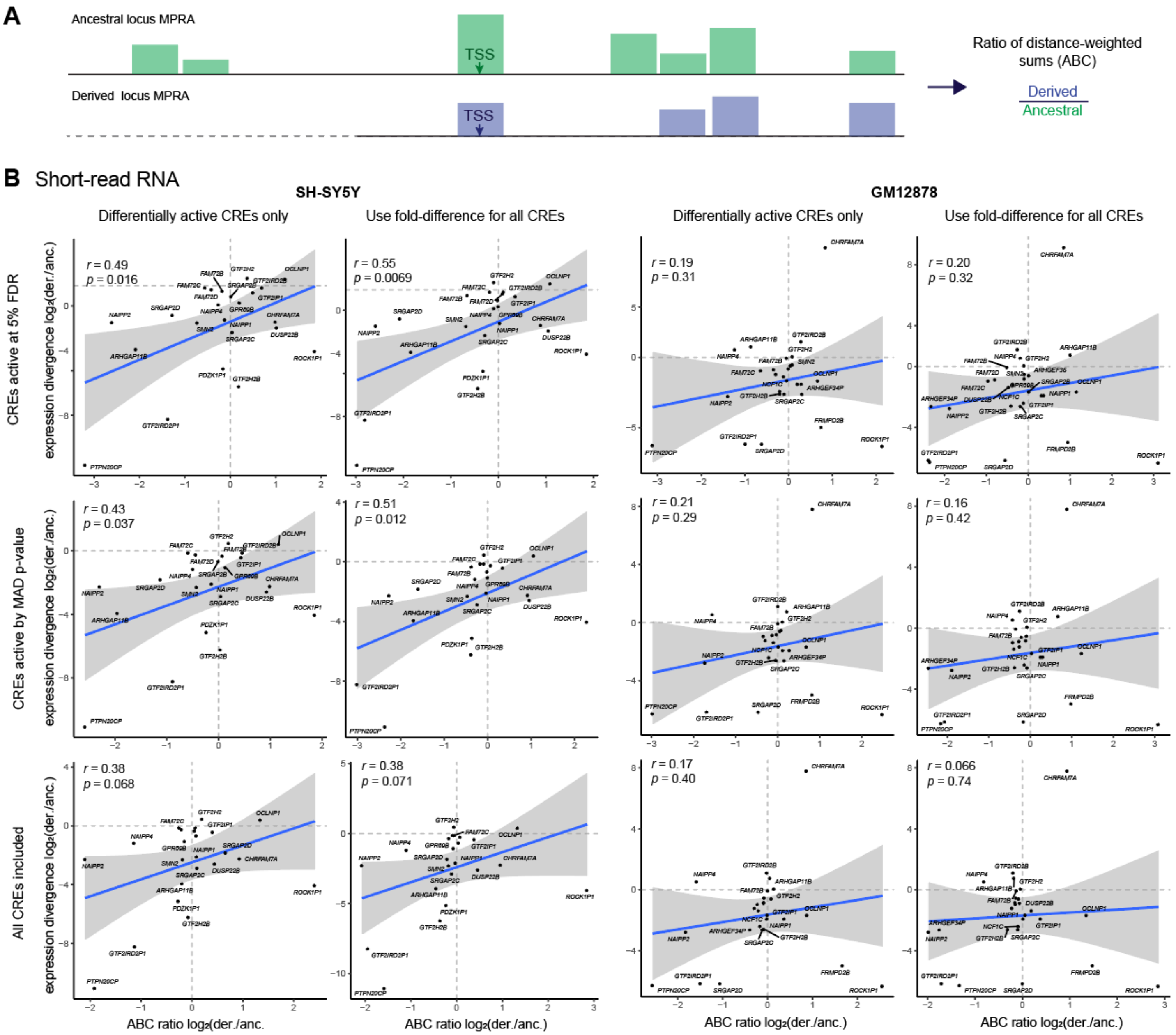
A
Ancestral locus MPRA
TSS
Derived locus MPRA
TSS
Ratio of distance-weighted sums (ABC)
Derived
Ancestral
B Short-read RNA
SH-SY5Y
GM12878
Differentially active CREs only
Use fold-difference for all CREs
Differentially active CREs only
Use fold-difference for all CREs
CREs active at 5% FDR
CREs active by MAD p-value
All CREs included
expression divergence log2(der./anc.)
r = 0.49
p = 0.016
r = 0.55
p = 0.0069
r = 0.19
p = 0.31
r = 0.20
p = 0.32
r = 0.43
p = 0.037
r = 0.51
p = 0.012
r = 0.21
p = 0.29
r = 0.16
p = 0.42
r = 0.38
p = 0.068
r = 0.38
p = 0.071
r = 0.17
p = 0.40
r = 0.066
p = 0.74
ABC ratio log2(der./anc.)

**C** Long-read RNA

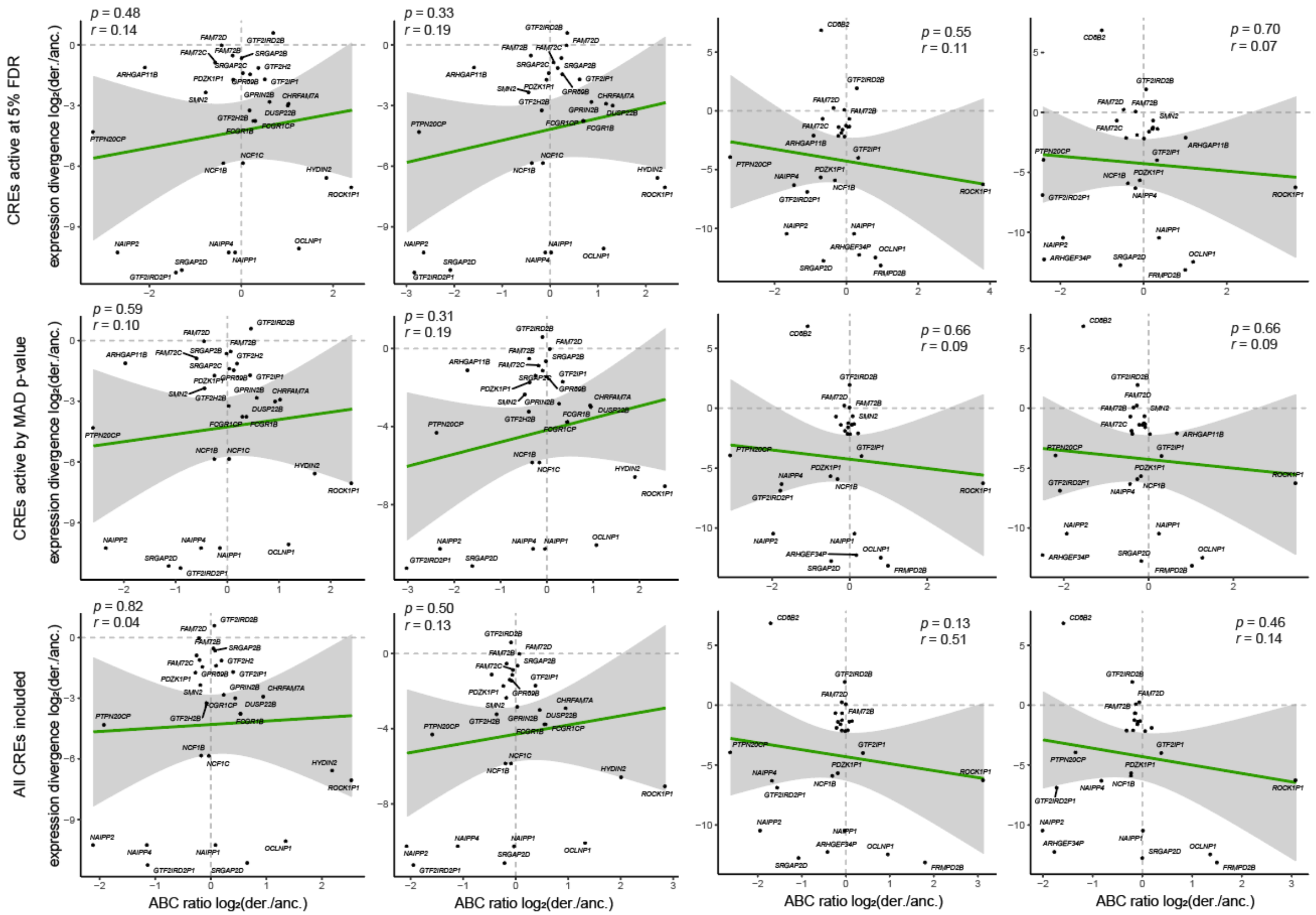


**Supplemental Fig. S14. ABC difference score evaluated with different sets of CREs. (A)** Short-read gene expression and **(B)** long-read (Kinnex) gene expression: For each cell type assayed by MPRA, mRNA expression divergence was correlated to the ratio of ABC scores for each derived-ancestral (der./anc.) gene pair. Each row shows decreasingly stringent definitions for CREs included, and each column shows a calculation using either differentially active CREs (5% FDR) or fold-difference values for all CREs regardless of significance. The Pearson's correlation and significance of the linear relationship is printed on each plot, and the 95% confidence interval for the regression is shaded in gray.

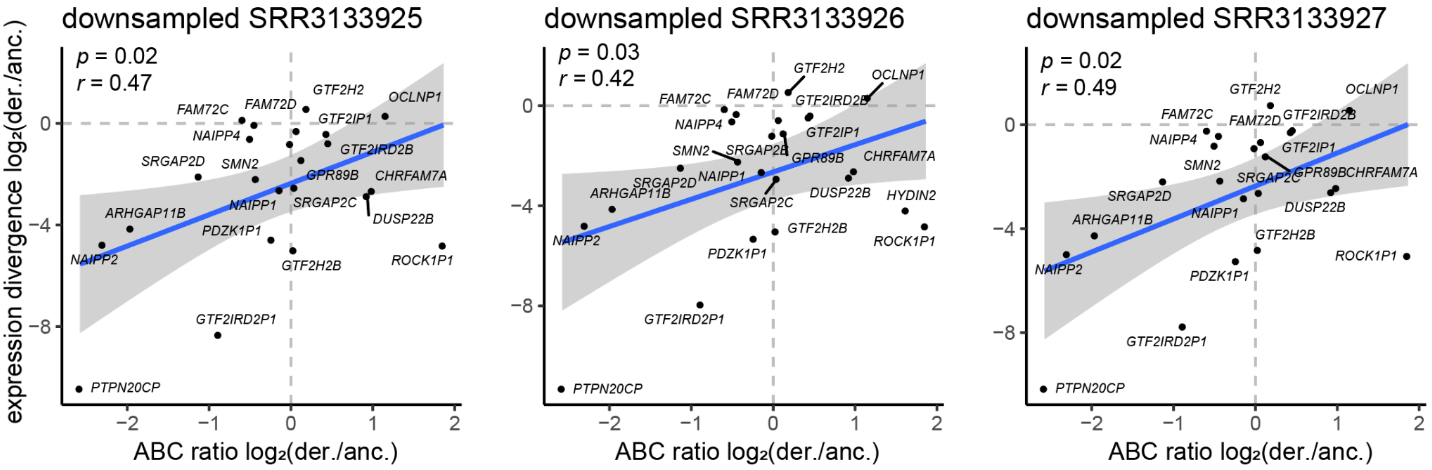


**Supplemental Fig. S15. ABC difference score evaluated on separate, downsampled replicates.** Correlation of ABC score ratios from the SH-SY5Y MPRA with expression divergence in SH-SY5Y (Pezzini et al., 2012). Each replicate was downsampled to match the read count in the Kinnex data. This comparison used differentially active CREs (5% FDR, at least one per pair active over control), as in Supplemental Fig. S14 (middle-left). Values shown are $\log_2$-tranformed ratios of derived (der.) to ancestral (anc.) expression. The Pearson's correlation and significance of the linear relationship is printed on each plot, and the 95% confidence interval for the regression is shaded in gray.

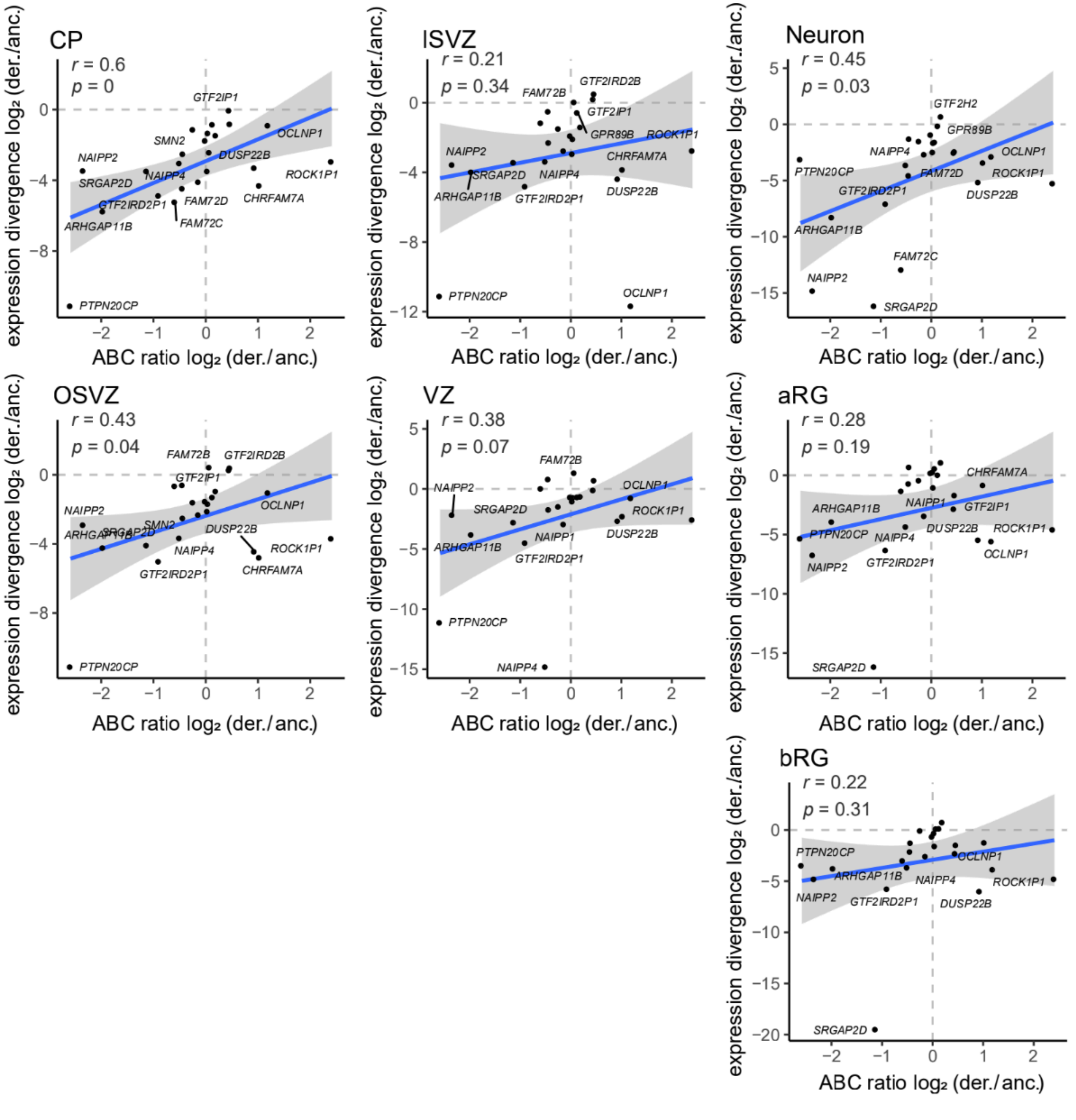


**Supplemental Fig. S16. ABC difference score evaluated with gene expression from primary brain.** Comparison of ABC score ratios from the SH-SY5Y MPRA with expression divergence from human fetal brain tissues (CP: cortical plate; iSVZ, inner subventricular zone; oSVZ, outer subventricular zone; VZ, ventricular zone; aRG, apical radial glia; bRG, basal radial glia; expression data from Florio *et al.*, 2015 and Fietz *et al.*, 2012). This comparison used differentially active CREs (5% FDR, at least one per pair active over control), as in Supplemental Fig. S14 (middle-left). Values shown are $log_2$-tranformed ratios of derived (der.) to ancestral (anc.) expression. The Pearson's correlation and significance of the linear relationship is printed on each plot, and the 95% confidence interval for the regression is shaded in gray.

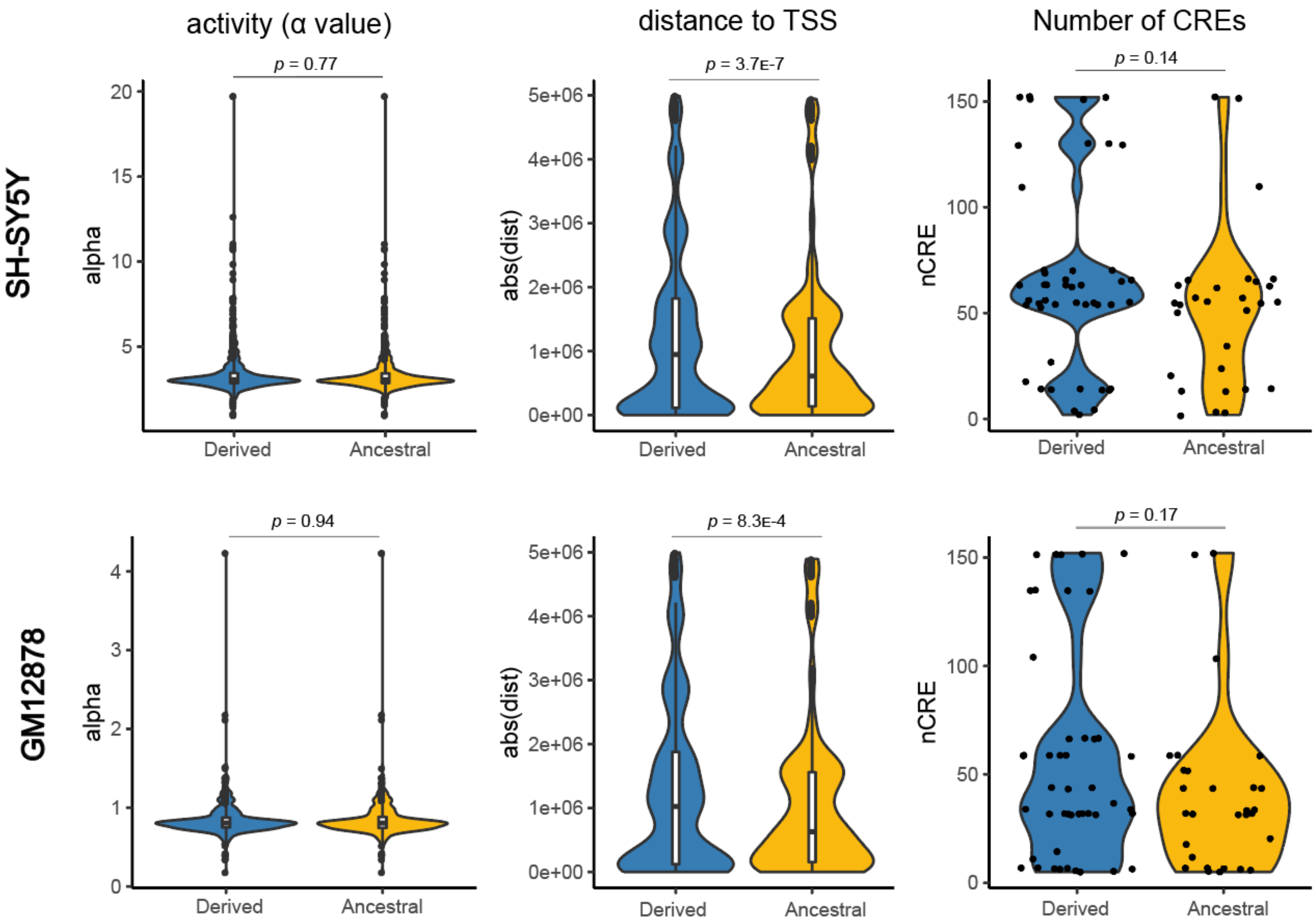


**Supplemental Fig. S17. Properties ancestral and derived CREs.** For MPRA performed in each cell type, the activity (alpha value) of each CRE, the distance between each CRE-TSS pair, and the number of CREs per gene used in the ABC score calculation are shown. Differences were assessed with a Wilcoxon rank sum test.

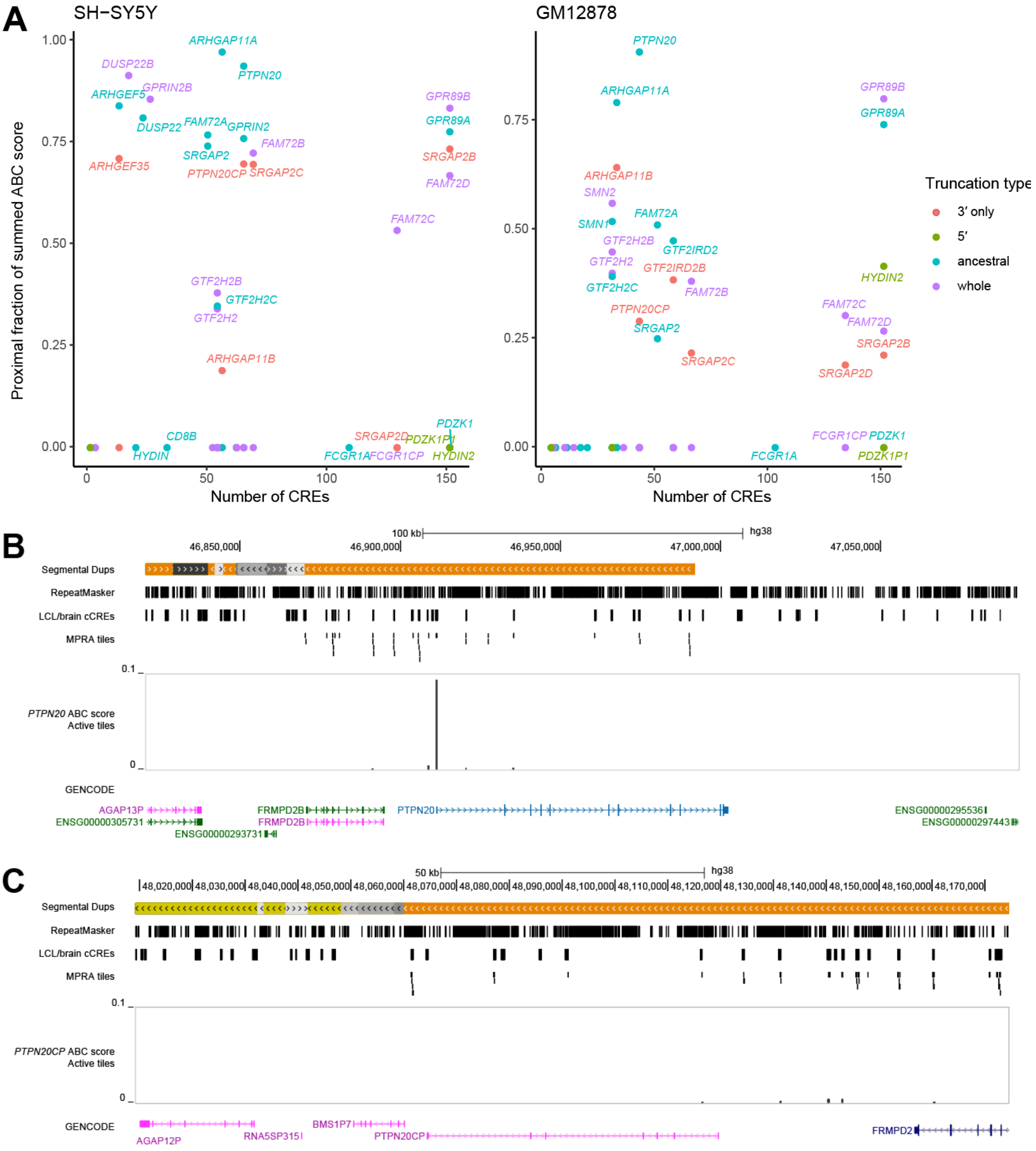
A
SH-SY5Y
GM12878
Proximal fraction of summed ABC score
Number of CREs
Truncation type
3′ only
5′
ancestral
whole
ARHGAP11A
PTPN20
DUSP22B
GPRIN2B
ARHGEF5
DUSP22
FAM72A
GPRIN2
FAM72B
SRGAP2
ARHGEF35
PTPN20CP
SRGAP2C
GPR89B
GPR89A
SRGAP2B
FAM72D
FAM72C
GTF2H2B
GTF2H2C
GTF2H2
ARHGAP11B
CD8B
HYDIN
SRGAP2D
PDZK1P1
PDZK1
FCGR1A
FCGR1CP
HYDIN2
SMN2
SMN1
GTF2IRD2
GTF2IRD2B
B
C
Segmental Dups
RepeatMasker
LCL/brain cCREs
MPRA tiles
PTPN20 ABC score
Active tiles
PTPN20CP ABC score
GENCODE
hg38
100 kb
50 kb
AGAP13P
ENSG00000305731
FRMPD2B
ENSG00000293731
ENSG00000295536
ENSG00000297443
AGAP12P
RNA5SP315
BMS1P7
FRMPD2

**D**

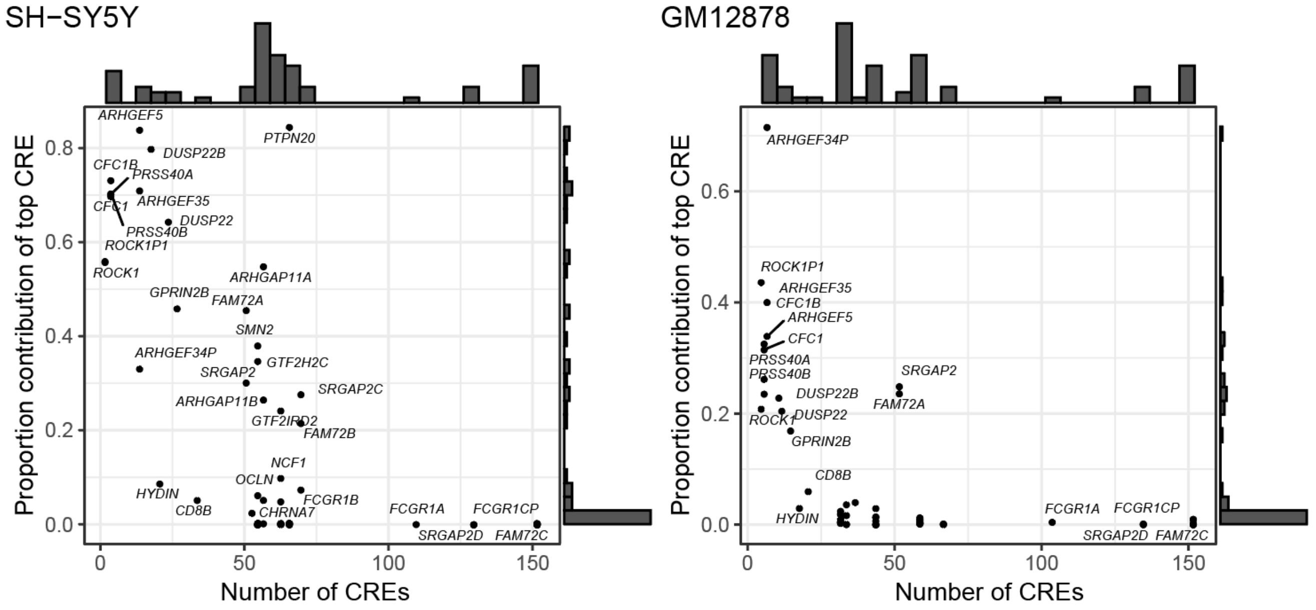


**Supplemental Fig. S18. Relative contributions of CREs to each gene's ABC score. (A)** All HSD genes assayed by MPRA are plotted by the number of CREs assigned to each gene during the ABC calculation and the proportional contribution of the of proximal CREs (within 1 kb) to that gene's score. Genes are colored by truncation status relative to the ancestral locus. **(B–C)** Visualization of divergent promoter activity of *PTPN20* and *PTPN20CP* in the UCSC Genome Browser. Tracks from top to bottom: Segmental duplications, with duplicated sequence >99% identity shown in orange; RepeatMasker annotation of transposable elements and simple repeats; ENCODE merged human LCL and brain CRE used to design MPRA; tested MPRA 200mer tiles; ABC scores (activity normalized by distance from TSS) for active elements used for paralog comparison; GENCODE annotation. **(D)** All HSD genes assayed by MPRA are plotted by the number of CREs assigned to each gene during the ABC calculation and the proportional contribution of the top CRE to that gene's score.

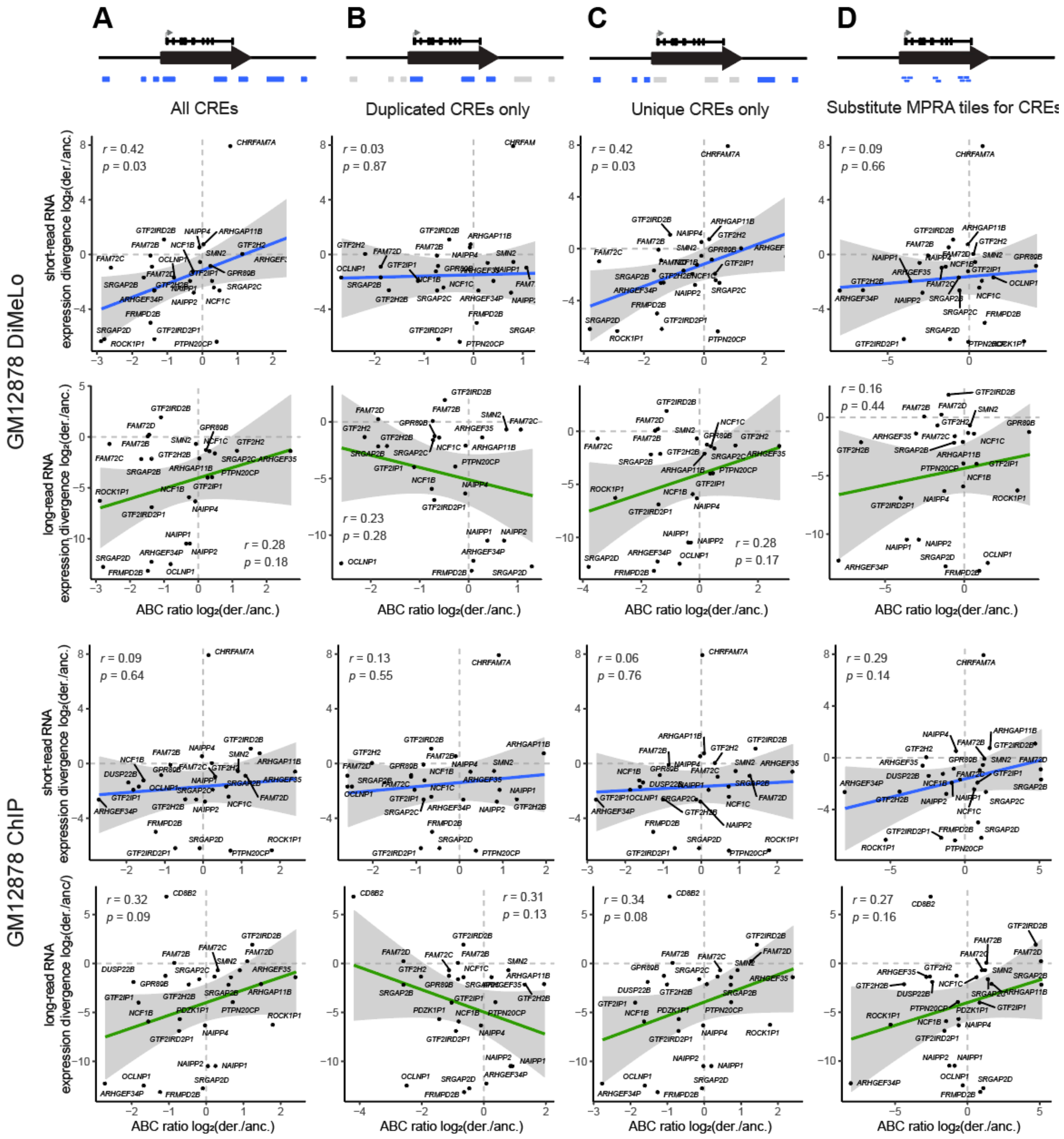


**Supplemental Fig. S19. Comparison of GM12878 expression divergence to ABC scores calculated from ChIP-seq and DiMeLo-seq.** For each gene expression dataset in GM12878, expression divergence was correlated to the ratio of summed ABC scores for each derived-ancestral (der./anc.) gene pair. ABC scores were calculated from **(A)** multi-mapped H3K27ac ChIP-seq and ATAC-seq and **(B)** H3K27ac signal from DiMeLo-seq. The comparison was performed using different sets of regions: (1) all CREs within 5 Mb, as defined by our ChIP-seq analysis; (2) CREs within HSDs; (3) CREs outside of HSDs; or (4) the exact positions tested by MPRA. The Pearson's correlation and significance of the linear relationship is printed on each plot. See Table S1 for a description of datasets used in this study.

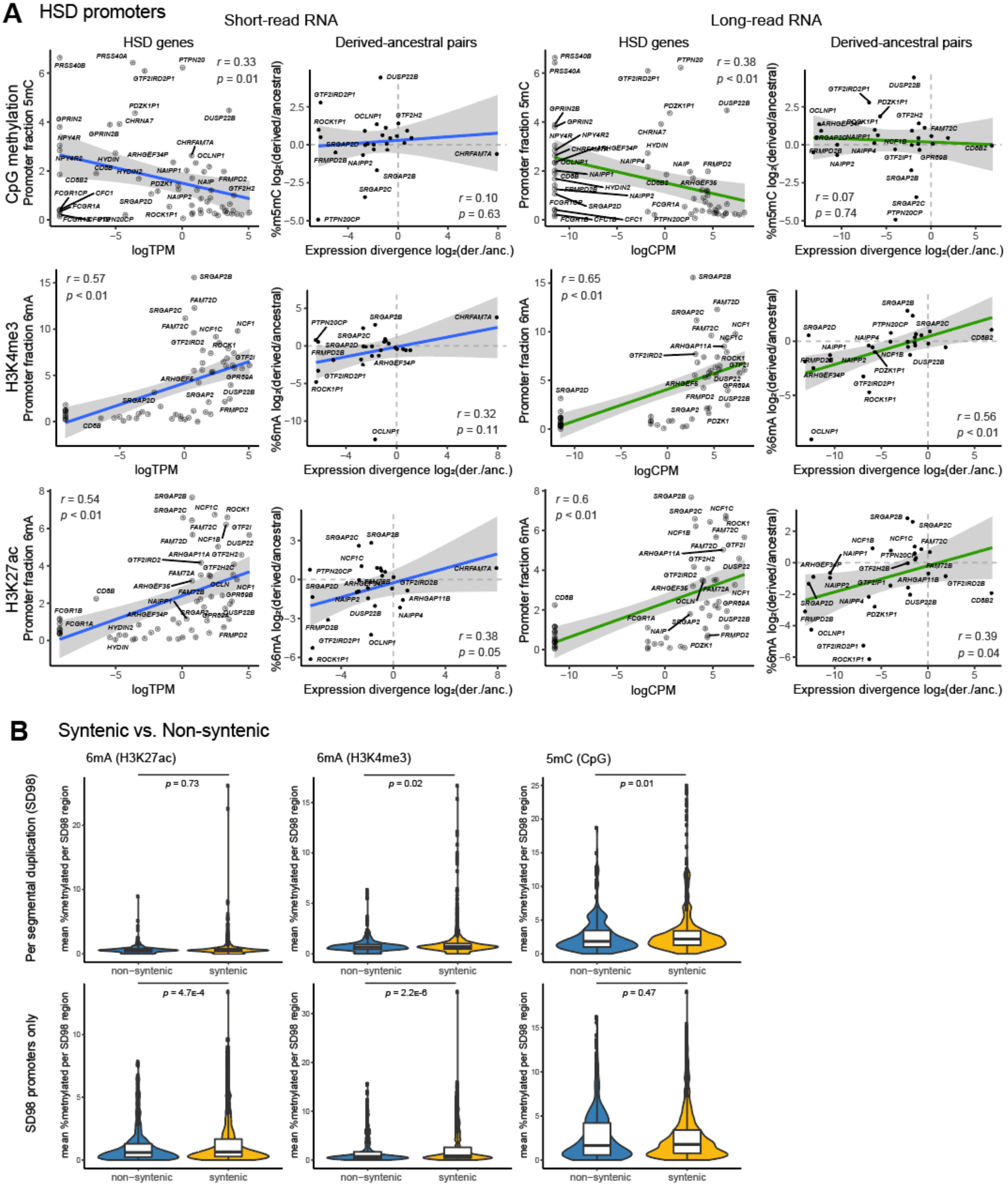
A
HSD promoters
Short-read RNA
Long-read RNA
HSD genes
Derived-ancestral pairs
CpG methylation
Promoter fraction 5mC
r = 0.33
p = 0.01
r = 0.10
p = 0.63
r = 0.38
p < 0.01
r = 0.07
p = 0.74
logTPM
logCPM
Expression divergence log2(der./anc.)
%m5mC log2(derived/ancestral)
H3K4me3
Promoter fraction 6mA
r = 0.57
p < 0.01
r = 0.32
p = 0.11
r = 0.65
p < 0.01
r = 0.56
p < 0.01
%6mA log2(derived/ancestral)
H3K27ac
r = 0.54
p < 0.01
r = 0.38
p = 0.05
r = 0.6
p < 0.01
r = 0.39
p = 0.04
B
Syntenic vs. Non-syntenic
6mA (H3K27ac)
6mA (H3K4me3)
5mC (CpG)
Per segmental duplication (SD98)
SD98 promoters only
mean %methylated per SD98 region
p = 0.73
p = 0.02
p = 0.01
p = 4.7E-4
p = 2.2E-6
p = 0.47
non-syntenic
syntenic

**Supplemental Fig. S20. Gene expression and CRE activity correlate with signal from long-read epigenome data from GM12878. (A)** For each expression dataset, correlations were calculated for all genes and all expressed ancestral-derived (anc./der.) gene pairs. Gene expression is plotted as log-transformed TPM or CPM, plus a pseudocount one order of magnitude below the smallest nonzero value (logTPM, logCPM), or expression divergence. Epigenomic signal was quantified from DiMeLo-seq (Maslan et al. 2024) as the fraction of modified bases for endogenous 5-methyl cytosine (5mC) at CpG sites or 6-methyl adenine (6mA, written by pA–Hia5 conjugated to anti-H3K4me3 or anti-H3K27ac). The Pearson's correlation and significance of the linear relationship is printed on each plot. **(B)** Global comparison of DiMeLo signal in segmental duplications with at least 98% sequence identity (SD98), previously classified as chimpanzee-syntenic or non-syntenic. The mean fraction methylated is calculated for each region (top) or promoter (bottom). Indicated *p*-values are from a Wilcoxon rank sum test.

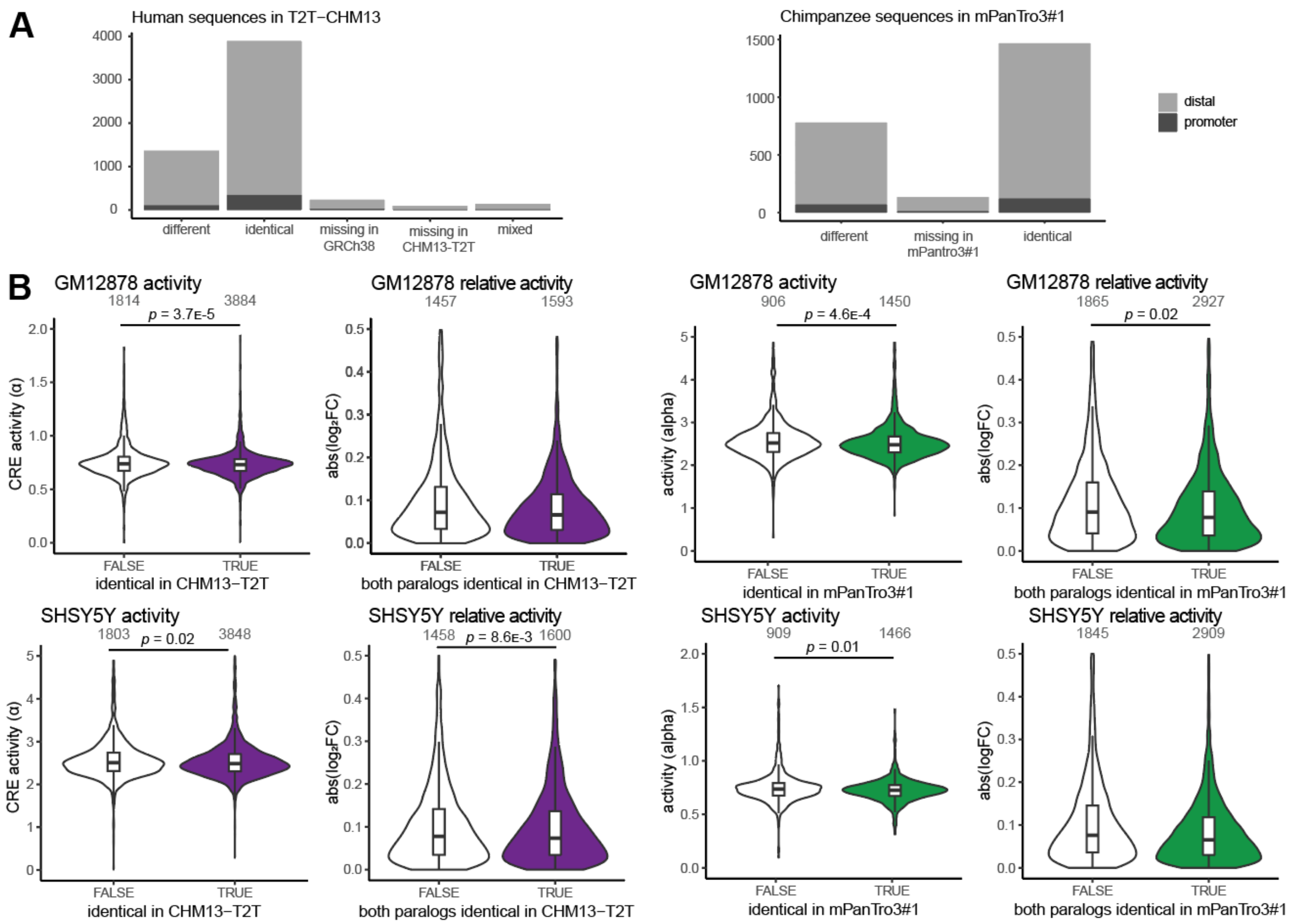


**Supplemental Fig. S21. MPRA results at sites differing in T2T assemblies.** Left: Human assembly comparison (GRCH38 and T2T-CHM13); right: chimpanzee assembly comparison (panTro6 and mPanTro3, haplotype 1). **(A)** MPRA tiles classified by whether they were identical between assemblies, failed to lift over, or had mixed classifications (paralogs which were identical in GRCh38). **(B)** MPRA results in GM12878 (top) and SH-SY5Y (bottom). Activity (alpha value) is plotted for all assayed human sequences, separated by whether the corresponding position in T2T-CHM13 or mPanTro3#1 was identical. Relative activity is plotted for all derived-ancestral gene pairs, separated by whether both sequences were identical between assemblies.
